# A survey of the mouse hindbrain in the fed and fasted state using single-nucleus RNA sequencing

**DOI:** 10.1101/2021.03.11.434948

**Authors:** Georgina K.C. Dowsett, Brian Y.H. Lam, John Tadross, Irene Cimino, Debra Rimmington, Anthony P. Coll, Joseph Polex-Wolf, Lotte Bjerre Knudsen, Charles Pyke, Giles S.H. Yeo

## Abstract

**Objective:** The area postrema (AP) and the nucleus tractus solitaris (NTS), located in the hindbrain, are key nuclei that sense and integrate peripheral nutritional signals and, consequently, regulate feeding behaviour. While single cell transcriptomics have been used in mice to reveal the gene expression profile and heterogeneity of key hypothalamic populations, similar in-depth studies have not yet been performed in the hindbrain.

**Methods:** Using single-nucleus RNA sequencing, we provide a detailed survey of 16,034 cells within the AP and NTS of the mouse, in the fed and fasted state.

**Results:** Of these, 8910 are neurons that group into 30 clusters, with 4289 coming from mice fed *ad libitum* and 4621 from overnight fasted mice. 7124 nuclei are from non-neuronal cells, including oligodendrocytes, astrocytes and microglia. Interestingly, we identified that the oligodendrocyte population was particularly transcriptionally sensitive to an overnight fast. The receptors GLP1R, GIPR, GFRAL and CALCR, which bind GLP1, GIP, GDF15 and amylin respectively, are all expressed in the hindbrain and are major targets for anti-obesity therapeutics. We characterise the transcriptomes of these four populations and show that their gene expression profiles are not dramatically altered by an overnight fast. Notably, we find that roughly half of cells that express GIPR are oligodendrocytes. Additionally, we profile POMC expressing neurons within the hindbrain and demonstrate that 84% of POMC neurons express either PCSK1, PSCK2 or both, implying that melanocortin peptides are likely produced by these neurons.

**Conclusion:** We provide a detailed single-cell level characterisation of AP and NTS cells expressing receptors for key anti-obesity drugs that are either already approved for human use or are in clinical trials. This resource will help delineate the mechanisms underlying the effectiveness of these compounds, and also prove useful in the continued search for other novel therapeutic targets.

## Introduction

Studies in humans and animal models have highlighted the central role of the brain in the control of appetite. Genetic and molecular approaches, for instance, have uncovered key circuits within the hypothalamus that sense and integrate peripheral nutritional signals, and as a consequence, regulate food intake and body weight. In addition, a number of nuclei residing within the hindbrain have also been identified as critical centres for the regulation of feeding behaviour. The area postrema (AP) is a circumventricular organ located within the medulla oblongata below the fourth ventricle, and is, like the hypothalamus, able to detect circulating chemical messengers in the blood and transduce them into neural signals and networks, relaying information to other brain areas [1]. Adjacent to the AP is the nucleus tractus solitaris (NTS) consisting of a collection of subnuclei which receive direct inputs from multiple locations including gustatory information from cranial nerves, and postprandial signals via vagal afferents [2], and thus acts as a visceral sensory relay station within the brain.

Both the AP and NTS have been implicated in the regulation of appetite [3], with a number of therapies for the treatment of obesity that are currently either in clinical trials or already on the market, targeting key receptors expressed within the one or both nuclei. For example, multiple long-acting GLP1R agonists are indicated for the treatment of Type 2 Diabetes, with liraglutide further indicated for the treatment of obesity, and semaglutide in submission for regulatory approval for obesity. These have been shown to cause weight loss in humans [4, 5] by acting on central GLP1R expressing neurons in both the hypothalamus and the NTS to suppress food intake [6–8]. The AP and NTS are also major sites of action for the hormone amylin, where it binds and activates Calcitonin receptors (CALCR) coupled to Ramp proteins [9]. Additionally, GDF15, a sentinel hormone that conveys states of somatic stress to the brain thus influencing appetite, acts exclusively through GFRAL receptors in the AP and NTS [10]. More recently, combinatorial therapies are being studied to assess the efficacy of targeting multiple circuits to enhance obesity treatments. Of interest, GLP1R agonists and dual amylin/calcitonin receptor agonists have been shown to have additive effects on reduction of food intake and body weight in rodents [11]. In addition, with activation of hypothalamic GIPR neurons resulting in a reduction of food intake [12], dual agonists of GLP1 and GIP receptors are being trialled as treatments for obesity and Type 2 Diabetes [for review see: 13]. Finally, a population of neurons that play a key role in the sensing of circulating nutritional signals and in so doing orchestrate appetite and peripheral metabolism, express Proopiomelanocortin (POMC). Within the brain, while POMC is primarily expressed in the arcuate nucleus of the hypothalamus (ARC), a smaller population exists in the NTS.

Multiple studies using single cell RNA sequencing have been performed to characterise neurons within the hypothalamus and comparing transcript expression across multiple nutritional states [14–17]. The transcriptional landscape of the AP upon treatment with amylin has been examined using so-called ‘bulk’ RNA-sequencing studies [9]. More recently single-nucleus RNA sequencing has been performed on 1,848 mouse AP neurons [18]. However, cells from the NTS were not included and only *ad libitum* fed mice were studied.

Here, using single-nucleus RNA sequencing (NucSeq), we perform a survey of 16,034 cells from the AP and NTS of the mouse, in the fed and fasted state. We report a detailed transcriptional profile of hindbrain neurons expressing GLP1R, GIPR, CALCR and GFRAL, and that their gene expression profiles are not dramatically altered by an overnight fast. Of note, we find that a large proportion of cells that express GIPR are not actually neurons, but oligodendrocytes. We also profile POMC expressing neurons within the hindbrain and demonstrate that the majority express PCSK1 and/or PCSK2, implying that melanocortin peptides are likely produced by these cells. Thus, we provide a detailed single-cell level characterization of neurons within the AP and NTS that express receptors for key anti-obesity drugs that are either already approved for human use or are in clinical trials.

## Materials and Methods

Mouse studies were performed in accordance with UK Home Office Legislation regulated under the Animals (Scientific Procedures) Act 1986 Amendment, Regulations 2012, following ethical review by the University of Cambridge Animal Welfare and Ethical Review Body (AWERB). All mice were obtained from Charles River. Twelve 6-8 week old C57BL/6J male mice were maintained in a 12-hour light/12-hour dark cycle (lights on 0700–1900), temperature-controlled (22°C) facility, with ad libitum access to food (RM3(E) Expanded chow, Special Diets Services, UK) and water. The day before sampling, 6 mice were subjected to an overnight fast for 16 hours.

### Nucleus dissociation

Mice were sacrificed by cervical dislocation and a portion of the hindbrain containing the AP and NTS was dissected and snap frozen on dry ice. Samples were stored at −80°C overnight. Samples from the same nutritional condition were pooled together to give 1 *ad libitum* sample and 1 fasted sample. The next day, homogenates were made using a dounce homogenizer in 1 ml homogenate buffer (100 μM DTT [Sigma, MO, Illinois, USA], 0.1% Triton X-100 [Sigma], 2X EDTA Protease Inhibitor [Roche, Basel, Switzerland], 0.4U/μl RNasin RNase inhibitor [Promega, Madison, WI, USA, 10000U, 40U/ml], 0.2U/μl Superase.In RNase Inhibitor [Ambion, Austin, TX, USA, 10000U, 20U/μl] in nuclei isolation medium [250 mM sucrose, 25 mM KCl (Ambion), 5mM MgCl_2_ (Ambion), 10 mM Tris buffer, pH 8.0 (Ambion) in nuclease free water (Ambion)] with 1μl/ml DRAQ5 [Biostatus, Loughborough, UK]) on ice. Homogenates were centrifuged at 900 x g for 10 minutes at 4°C and the supernatant discarded. The pellets were then resuspended in homogenate buffer with 25% Optiprep diluted in iodixanol dilution medium (250 mM sucrose, 150 mM KCl, 30 mM MgCl2, 60 mM Tris buffer, pH 8 in nuclease free water). Each suspension was layered on top of separate 29% Optiprep solutions and centrifuged for 20 minutes at 13,500 x g at 4°C. For each sample, the pellet was removed and resuspended in 1 ml of wash buffer (1% BSA, 0.4U/μl RNasin, 0.2U/μl Superase.In in PBS [Sigma]). Samples were passed through a 40μm cell strainer along with 0.7ml wash buffer.

### Fluorescent activated nucleus sorting

Fluorescent-activated cell sorting (FACS) was performed on both samples using the BD Influx cell sorter (Franklin Lakes, NJ, USA). The gating was set according to size and granularity using FSC and SSC to capture singlets and remove debris, and fluorescence at 647/670 nm to detect DraQ5 staining. Each sample was sorted into two separate tubes, with a total of 15,000 particles sorted into each tube containing 10μl wash buffer.

### Single nucleus RNA-sequencing

Sequencing libraries for the 4 (2 *ad libitum* and 2 fasted) single nuclei suspension samples were generated using 10X Genomics Chromium Single Cell 3’ Reagent kits (Pleasanton, CA, USA, Version 3) according to the standardised protocol. Briefly, nuclear suspensions were loaded on to the Chromium chip along with Gel Beads, Partitioning oil and master mix to generate GEMs containing free RNA. RNA from lysed nuclei was reverse transcribed and cDNA was PCR amplified for 19 cycles. The amplified cDNA was used to generate a barcoded 3’library according to manufacturer protocol and paired end sequencing was performed using an Illumina NovaSeq 6000 (San Diego, CA, USA, Read 1: 28bp, Read 2: 91bp). Library preparation and the sequencing was performed by the Genomics Core, Cancer Research UK Cambridge Institute.

### Single cell clustering and differential gene expression analysis

Reads were aligned to mouse genome (GRCm38) using the CellRanger package version 4.0 (10X Genomics). Downstream analysis on the raw count matrices were performed using the Seurat package v 3.1.1[19, 20]. Briefly nuclei expressing less than 200 features, or less than 1000 transcripts were removed as low-quality reads. Additionally, nuclei with more than 6000 different features were removed as these were likely doublets. The data was log-normalised and scaled prior to PCA, followed by unsupervised clustering analysis using the Louvain algorithm and non-linear dimensional reduction via T-distributed Stochastic Neighbour Embedding (tSNE). Any clusters that showed high expression of more than one cell type marker (e.g. neuron, astrocyte) were further removed. Marker genes for each cluster were identified using Wilcoxon Ranked Sum test and receiver-operating curve (ROC) analysis. Additionally, differential expression of genes between nuclei from *ad libitum* fed and overnight fasted mice within each cluster was analysed using Wilcoxon Ranked Sum test with adjusted p-values based on Bonferroni correction using all the features within the dataset.

To characterise populations of GLP1R, GFRAL, CALCR, GIPR and POMC individually, subsets of nuclei expressing at least 1 raw UMI count of the gene of interest were created. If the gene was found to be expressed in mainly the neurons then the subset was taken from the neuronal dataset (this was the case for GLP1R, CALCR and GFRAL), however if there was widespread transcript expression within the non-neuronal clusters, the subset was taken from the whole dataset (POMC and GIPR). Subsets were re-clustered according to the Seurat pipeline (see above). Marker genes for each cluster were identified using Wilcoxon Ranked Sum test and ROC analysis. Violin plots were generated using the Seurat package. The differential gene expression cut off used was unadjusted p<0.05 unless otherwise stated.

To characterise receptor and peptide expression profiles of these subsets of nuclei, any gene that was expressed in a minimum of 50% of cells in one of the clusters was kept for analysis. The mean relative expression level of each gene was calculated for each cluster to characterise the expression profiles of these cells. To determine the differential gene expression of these subsets in response to a fast, the genes that were used for expression profiling were analysed for changes in gene expression within each cluster between fed and fasted nuclei using the EdgeR package [21]. For receptors and transporters, gene type was determined based on the classification from Ingenuity Pathway Analysis (Qiagen). For peptide hormones, genes annotated with the gene ontology (GO) term: GO:0005179 (hormone activity) were included.

In addition, genes whose roles in neuronal transmission are well established (*Slc17a6*, *Slc17a7, Slc32a1, Gad1, Gad2, Slc6a5, Ddc, Dbh, Th, Slc6a3, Tph1, Slc6a4, Chat, Slc18a3)* and expressed in a minimum of 10% of cells from one cluster in each subset were also analysed in the same way (average expression level and differential gene expression). The same was done for a number of oligodendrocyte markers for the GIPR and POMC subsets as these had oligodendrocyte clusters (*Olig1, Olig2, Olig3, Cldn11, Mbp, Mog, Sox10, Pdgfra, Cspg4*).

### Pathway analysis

Ingenuity pathway analysis (IPA; Qiagen) was used to identify pathways that were either upregulated or downregulated between the fed and fasted condition.

## Results

### Single nucleus RNA-sequencing survey of 16,034 nuclei from the mouse AP and NTS

We isolated and performed NucSeq on a total of 16,034 nuclei from the AP/NTS of 12 male mice, six fed *ad libitum* and six fasted overnight. A median of 2419 different transcripts were detected per individual nucleus (**Figure 1a**). Unsupervised clustering analyses (see Methods) separated the nuclei into 41 different clusters on a tSNE plot (**Figure 1a**), with the cell type of each cluster determined based on the expression of canonical markers (**Figure 1c**).

**Figure 1.**
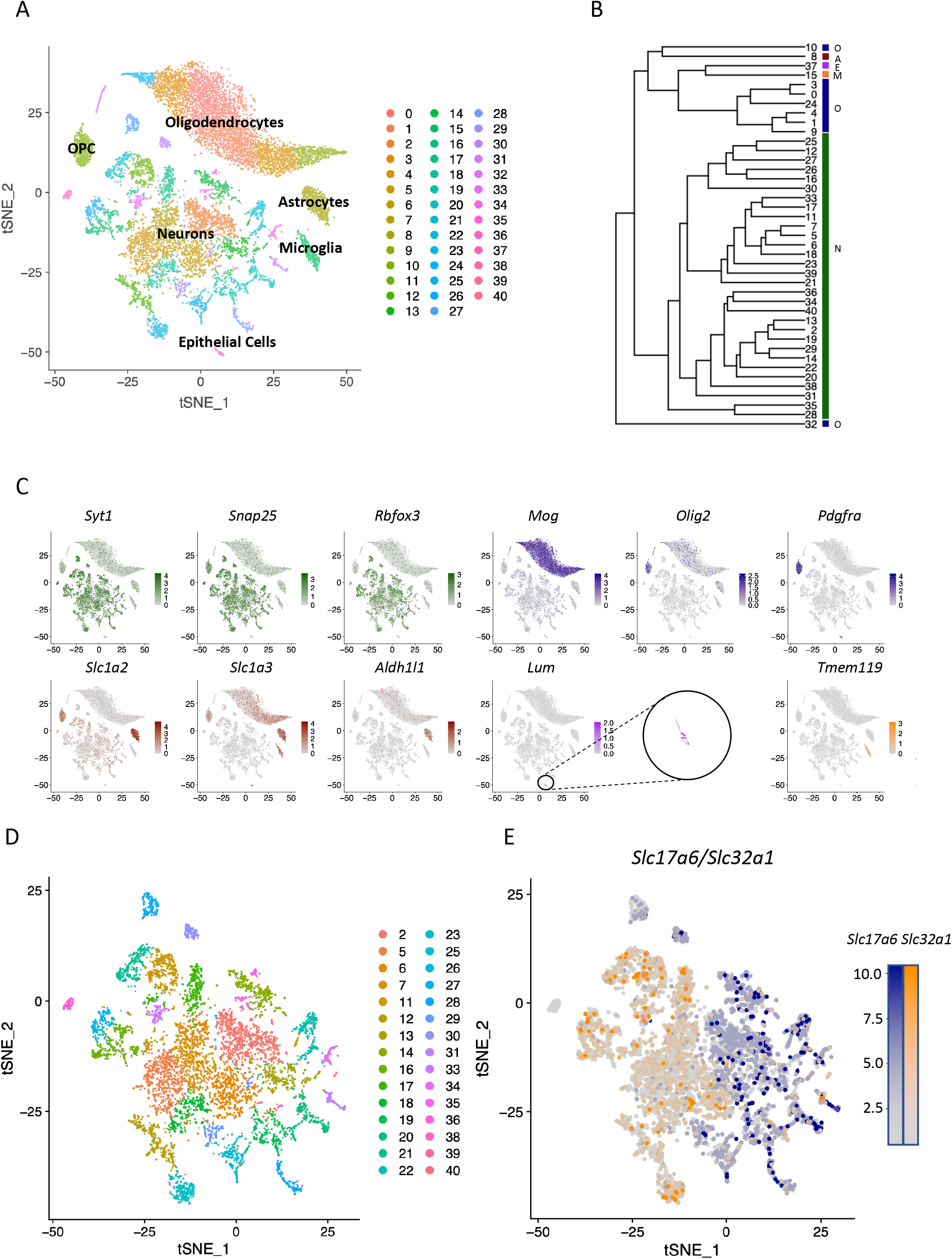
Single-nucleus RNA sequencing of the AP and NTS reveals 41 populations. (A) tSNE plot showing 41 clusters including 8 Oligodendrocyte and Oligodendrocyte Precursor Cells (OPC) clusters, 1 Astrocyte cluster, 1 microglia cluster, 1 epithelial cell cluster and 30 neuronal clusters. (B) Phylogenetic tree showing the hierarchy of the clusters. Similar cell clusters are grouped together. (Abbreviations: O = Oligodendrocytes, A = Astrocytes, E = Epithelial cells, M = Microglia, N = Neurons). (C) Feature plots displaying the relative expression levels of cell type markers for neurons (green, top left), Oligodendrocytes and OPC’s (blue, top right), astrocytes (red, bottom left), epithelial cells (pink, bottom centre) and microglia (orange, bottom right). (D) tSNE plot showing the 30 neuronal clusters (8910 nuclei). Cluster numbers remain unchanged from figure 1A. (E) Scaled expression levels of *Slc17a6* (blue) and *Slc32a1* (orange) in neuronal nuclei displayed as a combined feature plot.

11 of the 41 clusters encompassed 7124 non-neuronal cells (**Figure 1b**). A large population of oligodendrocytes, by marked expression of *Mog*, accounted for seven clusters that are closely adjacent to each other, with maturity of the oligodendrocytes, as marked by a corresponding decrease in expression levels of *Olig2*, increasing from left to the right. There was also a cluster of oligodendrocyte precursor cells (OPCs) marked by their expression of *Pdgfra*. A single cluster of astrocytes was identified by expression of the excitatory amino acid transporters 1 and 2 (*Slc1a3* and *Slc1a2*), as well as *Aldh1l1*. Additionally, there was one cluster of microglia (marked by expression of *Tmem119*) and a cluster of epithelial cells (expressing *Lum*) (**Figure 1c**).

The remaining 30 clusters were formed from 8910 neurons, all of which were characterised by their high relative expression of *Syt1*, *Snap25* and *Rbfox3* (**Figure 1c**). We focused on these 30 neuronal clusters (**Figure 1d**) and performed differential gene expression analysis to identify their key transcriptional markers (summarized in **Figure S1**). The primary driver differentiating the transcriptional profile of the neurons was their expression of either *Slc17a6* (vGlut2) and *Slc32a1* (vGat), which are excitatory and inhibitory markers respectively (**Figure 1e**). There were two clusters with low expression of both *Slc17a6* and *Slc32a1*; Cluster 36, which additionally expressed the cholinergic neuronal markers *Slc5a7* (encoding the high affinity choline transporter) and *Chat* (Choline acetyltransferase) (**Figure S1**); and cluster 27, that additionally expressed *Slc17a7* (vGlut1), *Ngf* and *Fam107b* (**Figure 1e**; **Figure S1**). Other notable populations include cluster 26, which is marked by the expression of *Npy*; clusters 31 and 34, that both express *Ddc* (DOPA decarboxylase; converting L-DOPA into dopamine) and *Dbh* (Dopamine-β-hydroxylase; which converts dopamine into noradrenaline), implying that these are noradrenergic neurons; and cluster 40, which is likely to be serotonergic due to expression of *Tph2* (Tryptophan Hydroxylase 2) and *Slc6a4* (a serotonin transporter) (**Figure S1**).

### The transcriptomic response of AP and NTS cell populations to an overnight fast

We next examined the consequences of an overnight fast on the transcriptional profile of AP and NTS cells, with 3378 non-neuronal and 4289 neuronal nuclei (total 7667) profiled from fed mice; and 3746 non-neuronal and 4621 neuronal nuclei (total 8367) profiled from fasted mice (**Figure 2**). Most of the neuronal clusters displayed a relatively equal number of fed:fasted nuclei, the exceptions being clusters 6 (markers *Cdh8* and *Sox2ot*), 25 (*Zic4* and *Grin2c*) and 40 (*Slc6a4* and *Tph2*), which had disproportionate number of nuclei from *ad libitum* fed mice (84%, 67.8% and 78.3% respectively); and clusters 13 (*Tac1* and *Dlk1*) and 36 (*Slc5a7* and *Chat*) that had an overrepresentation of nuclei from fasted mice (66.1% and 69.1% respectively) (**Figure 2b**).

**Figure 2.**
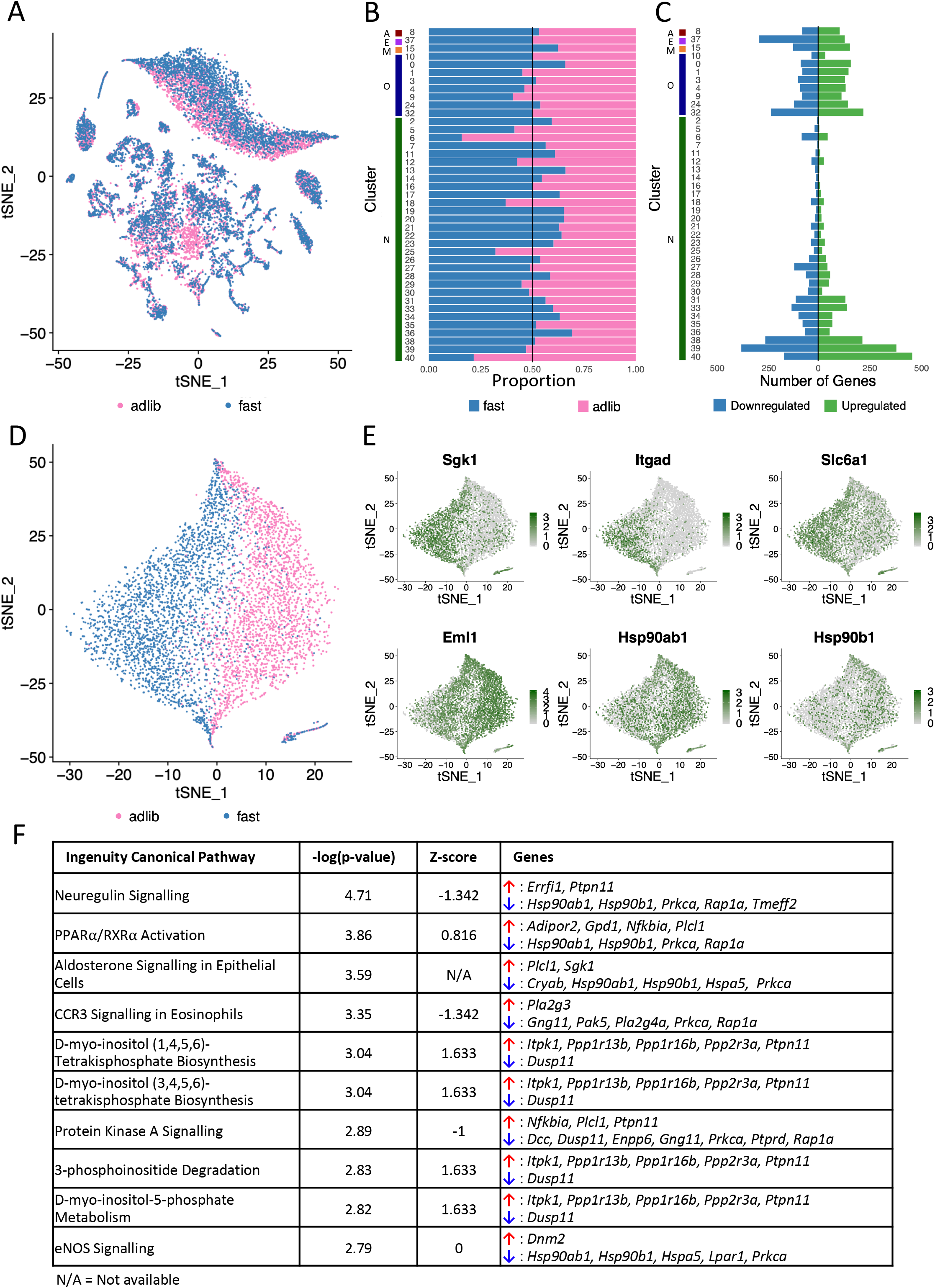
The effect of nutritional state on transcriptomic expression in mouse AP/NTS cells. (A) tSNE plot revealing the nutritional status of each cell within the dataset. A total of 7667 nuclei came from mice fed *ad libitum* (pink), and 8367 cells came from mice who were fasted overnight (blue). (B) Graph showing the proportion of nuclei within each cluster that originated from mice fed *ad libitum* (pink) or overnight fasted mice (blue). (C) Graph showing the number of genes upregulated (green) and downregulated (blue) in response to an overnight fast in each cluster. Genes included were significantly differentially regulated (P<0.05). (B-C) Cluster numbers listed down the left-hand side and the cell types of each cluster identified: N = neurons, O = Oligodendrocytes and OPCs, M = Microglia, E = Epithelial cells, A = Astrocytes. (D) A tSNE displaying a subset of oligodendrocyte nuclei (not including the OPC cluster), revealing the nutritional status of each cell (*ad libitum* in pink, fasted in blue). (E) Relative expression levels of top differentially expressed genes within the oligodendrocyte subset, displayed as feature plots. (F) Top 10 canonical pathways affected by an overnight fast in the oligodendrocyte population. A negative z-score represents an overall downregulation of the pathway, and a positive z score represents an overall upregulation of the pathway in the fasted state. Genes involved in each pathway whose expression was differentially regulated were highlighted for each pathway.

The population of oligodendrocytes were surprisingly transcriptionally responsive to an overnight fast (**Figure 2c**). Fasting, in fact, turned out to be the main driver of the oligodendrocyte clustering (**Figure 2d**). The oligodendrocytes showed consistently high levels of differential gene expression in the fasted state, with 6 out of the 7 clusters exhibiting more upregulated than downregulated genes (**Figure 2c**). When the entire oligodendrocyte population was considered as a whole, the top differentially regulated genes in response to fasting included upregulation of *Sgk1* (Serum/Glucocorticoid Regulated Kinase 1)*, Itgad* (Integrin alpha-D) encoding the alpha subunit of an integrin glycoprotein, and the GABA transporter 1 gene *Slc6a1,* which transports GABA into the cell. Additionally, downregulated genes included *Eml1* (Echinoderm microtubule associated protein-like 1), and two heat shock proteins *Hsp90ab1* and *Hsp90b1* (**Figure 2e**). Pathway analysis revealed that neuregulin signalling, important in myelination which involves *Hsp90ab1* and *Hsp90b1*, was downregulated (**Figure 2f**)[22]. Furthermore, a number of pathways involved in inositol metabolism, thought to regulate oligodendrocyte differentiation and survival [23, 24], and PPARα/RXRα signalling were upregulated in the fasted state (**Figure 2f**). Interestingly, expression of 25 of the differentially expressed genes in oligodendrocytes are regulated by the transcription factor *Tcf7l2*, including *Sgk1, Ptpn11, Rap1a* and *Ppp1r16b* all of which are implicated in the neuregulin or inositol biosynthesis pathways (**Figure 2f**).

### Characterisation of neurons expressing key anti-obesity therapeutic targets in AP and NTS

Next, we focussed on detailed analyses of four different populations of neurons that express the receptors GLP1R, GIPR, GFRAL and CALCR, which bind GLP1, GIP, GDF15 and amylin respectively, and are major targets for anti-obesity therapeutics.

#### GLP1R

We identified a total of 173 *Glp1r* expressing nuclei in 24 of the 30 neuronal clusters (**Figure 3a**). When we extracted and analysed these neurons in isolation, they fell into three clusters (**Figure 3b**). Cluster 0, the largest of the three, was characterised by the expression of *Kctd16* and *Alcam*, cluster 1 was marked by *Ddc*, and cluster 2 by *Slc17a7* (**Figure 3c**).

**Figure 3.**
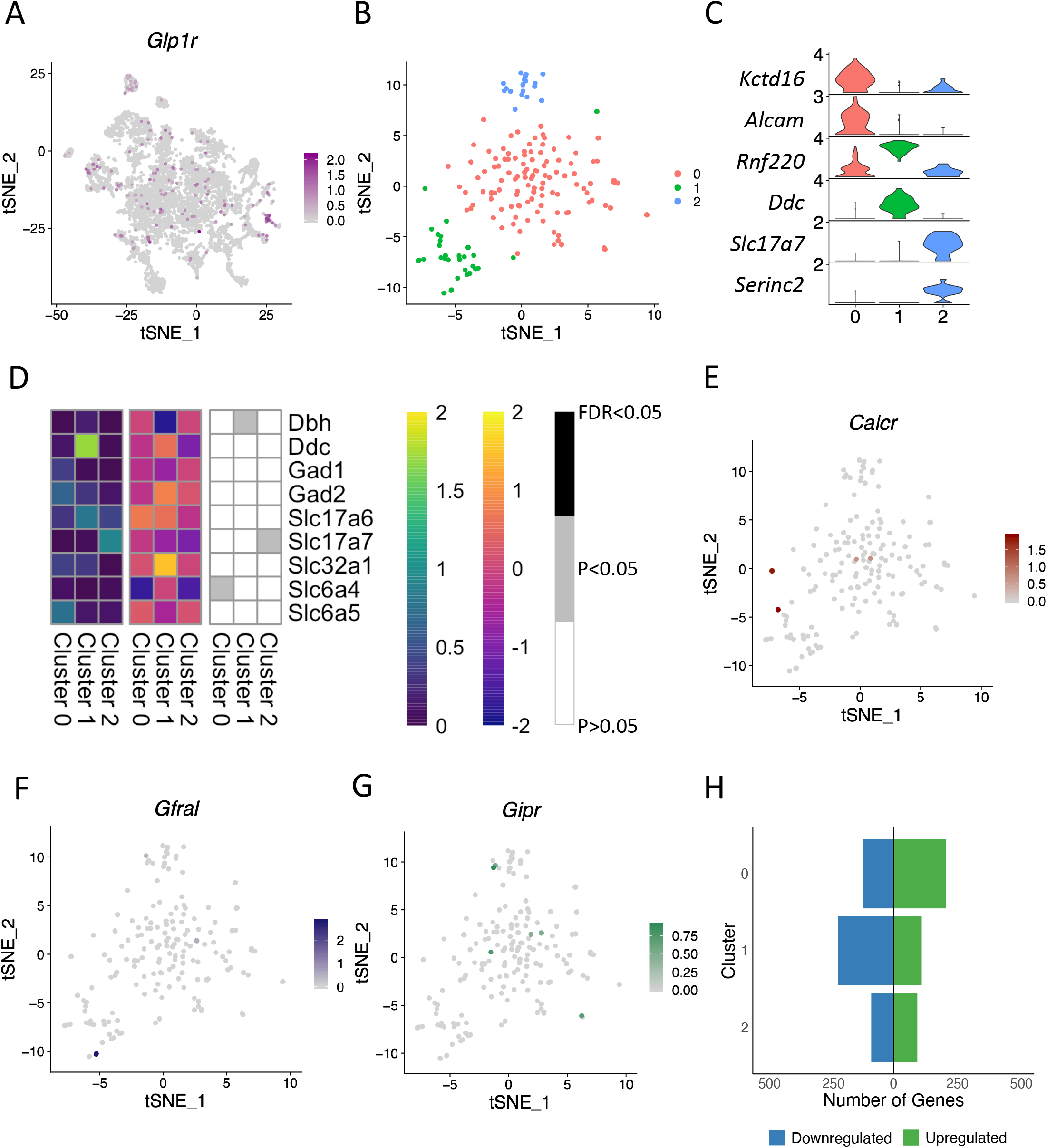
Characterisation of GLP1R expression in AP and NTS neurons. (A) Feature plot revealing relative expression levels of *Glp1r* in the 30 neuronal clusters. (B) tSNE plot of the 173 GLP1R neurons, grouped into 3 clusters. (C) Violin plots showing relative expression levels of 2 marker genes for each cluster of the GLP1R subset. The highest relative expression value for each gene is displayed on the left of the plot. (D) Heatmaps profiling the expression of canonical neuronal markers in each GLP1R cluster. Left: Average relative expression level in each cluster; Middle: Differential expression of each gene per cluster (Log2FC values); Right: Determining whether gene expression was significantly changed in the fasted state. (E-G) Feature plots showing relative expression levels of *Calcr* (E), *Gfral* (F), and *Gipr* (G) in the GLP1R subset. (H) Graph showing the number of genes upregulated (green) and downregulated (blue) in response to an overnight fast in each cluster. Genes included were significantly differentially regulated (P<0.05).

Cluster 1 consisted of both excitatory and inhibitory neurons, with 48.4% expressing *Slc17a6* and 22.6% expressing *Slc32a1* (**Figure 3d**). In cluster 2, 83.3% of nuclei expressed *Slc17a7* (vGlut1) and 38.9% expressed *Slc17a6* (vGlut2, **Figure 3d**). Cluster 0 neurons predominantly expressed inhibitory neuronal markers, with 46% of nuclei expressing *Gad2* and *Slc6a5* (average relative expression levels: 0.64 and 0.75 respectively). These data show that GLP1R neurons within the mouse hindbrain can either be excitatory or inhibitory. Next, we investigated the expression of *Gfral, Gipr* and *Calcr* in the *Glp1r* subset. These four populations of neurons appear largely distinct, with only 6 *Glp1r* nuclei co-expressed *Gipr*, 5 co-expressed *Calcr* and 4 co-expressed *Gfral* (**Figure 3e-g**), and could suggest that agonists to all these receptors act largely independently of each other. However, while this experiment is designed to identify the presence of transcripts it is entirely possible that cells expressing low levels of transcripts could be missed. A detailed expression profile of GLP1R neurons is provided in **Figure S2**.

To determine whether GLP1R neurons are sensitive to an overnight fast, we performed differential gene expression analysis in all 3 clusters (See methods). Clusters 0 and 1 showed the highest changes in gene expression in the fasted state (326 and 327 differentially expressed genes respectively; **Figure 3h; Table S1**). Notably, in cluster 0, *Ptgds* expression was significantly upregulated (LogFC = 1.1566, P = 6.40×10^−8^, False Discovery Rate [FDR] = 0.0003), and in cluster 1 *Oxr1* (Oxidation resistance 1) was significantly downregulated in response to an overnight fast (LogFC = −1.9133, P = 3.78×10^−6^, FDR = 0.0156). Pathway analysis revealed the synaptogenesis signalling pathway to be significantly changed in two of the clusters (downregulated in cluster 0, upregulated in cluster 2), and calcium signalling to be upregulated in cluster 2 (**Table S2**).

#### GIPRs are expressed on both neurons and oligodendrocytes in the mouse hindbrain

GIPR expression within the hypothalamus has been characterised using histological techniques in combination with single cell RNA-sequencing [12], however less is known about GIPR expressing cells within the hindbrain. We identified 436 *Gipr* expressing nuclei that were found in all but two clusters within the entire hindbrain dataset (**Figure 4a**). Neuronal cluster 21 contained the highest proportion of *Gipr* nuclei, with 11.5% of this cluster expressing *Gipr*. Marker genes for this cluster included *Ccbe1* and *Zeb2* (**Figure S1**). Unexpectedly, a large number of *Gipr*^+^ nuclei were oligodendrocytes (**Figure 4a**), with cluster 9 exhibiting the highest percentage of *Gipr* nuclei (13.4%).

**Figure 4.**
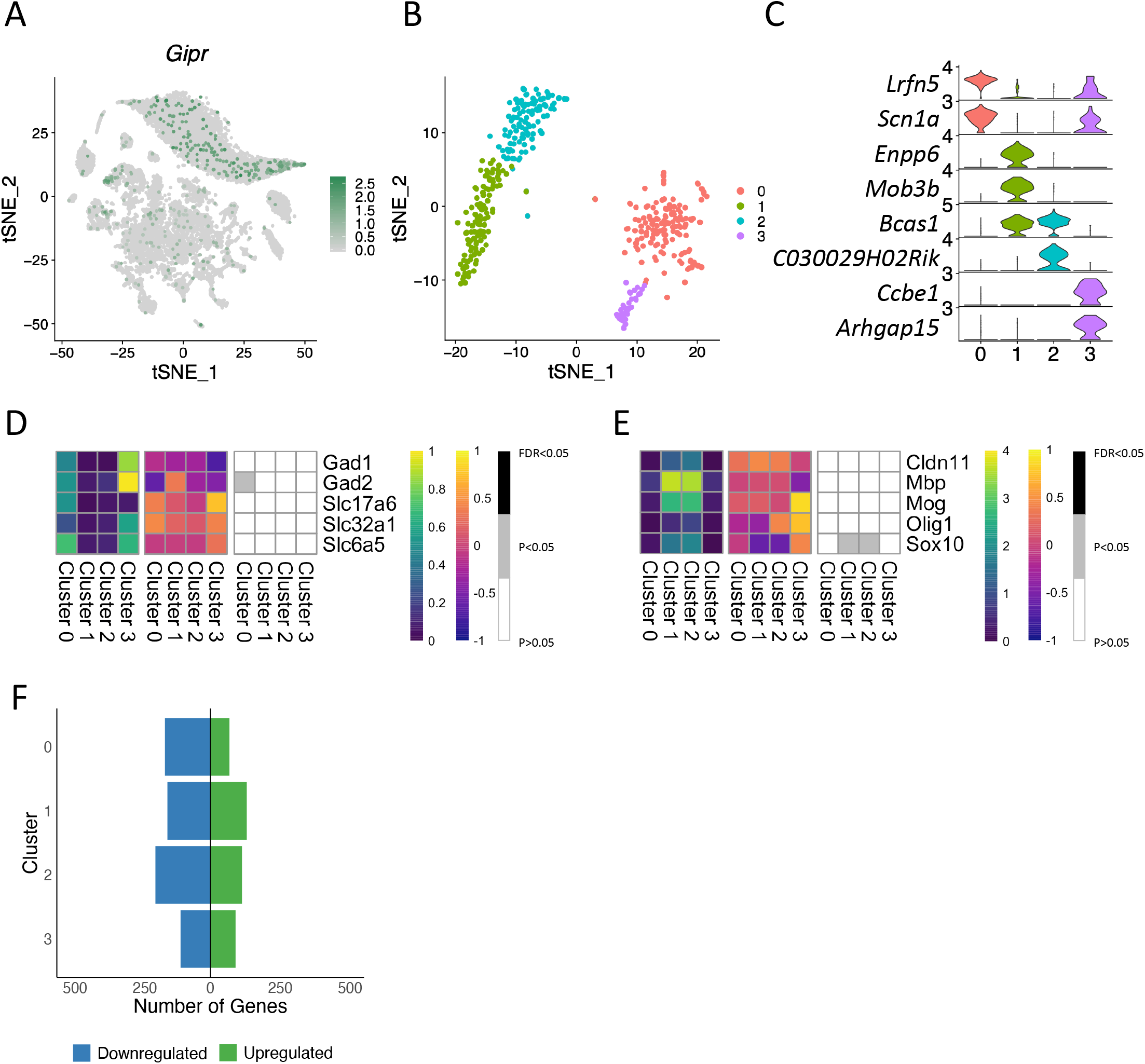
GIPR is expressed in neuronal and non-neuronal cells within the mouse AP and NTS. (A) Feature plot displaying the relative expression levels of *Gipr* in the overall dataset. (B) tSNE plot of the 436 GIPR cells, which grouped into 4 clusters. (C) Violin plots showing relative expression levels of 2 marker genes for each cluster of the GIPR subset. (D-E) Heatmaps profiling the expression of canonical neuronal (D) and oligodendrocyte (E) markers in each GIPR cluster. Left: Average relative expression level in each cluster; Middle: Differential expression of each gene per cluster (Log2FC values); Right: Identifying whether gene expression was significantly changed in the fasted state. (F) Graph showing the number of genes upregulated (green) and downregulated (blue) in response to an overnight fast in each cluster. Genes included were significantly differentially regulated (P<0.05).

When we re-clustered the 436 *Gipr* expressing nuclei, they formed 4 clusters, 2 of oligodendrocytes and 2 neuronal (**Figure 4b**). Both oligodendrocyte clusters expressed *Bcas1*, a myelin associated protein, as well as *Mbp* and *Mog* (Cluster 1: 100% and 99.2%; Cluster 2: 99.1% and 91.8% respectively). Additionally, oligodendrocyte cluster 1 expressed *Enpp6*, an enzyme involved in choline metabolism which is typically expressed in maturing oligodendrocytes [25]. Both clusters of neurons express *Scn1a* and *Lrfn5*, with cluster 3 expressing *Ccbe1* and *Arhgap15* (**Figure 4c**). 39.9% of cluster 0 expressed the glutamatergic marker *Slc17a6* (average relative expression: 0.37), and 39.9% express *Slc6a5* (average expression level of 0.45). Cluster 3 was predominantly GABAergic, with 48.7% and 61.5% expressing *Gad1* and *Gad2* (**Figure 4d**). Further characterisation of expression profiles of GIPR cells is found in **Figure S3.**

Out of the 436 nuclei, 190 came from fasted mice. Differential expression analysis in each cluster showed the highest changes in gene expression were within the oligodendrocyte clusters, with 284 and 310 differentially expressed genes in response to a fast in clusters 1 and 2 respectively (**Figure 4f; Table S3**). Similar to GLP1R neurons (**Table S1**), *Ptgds* was also significantly upregulated in cluster 0 and cluster 2 (LogFC = 1.4381, P = 3.14×10^−10^, FDR = 8.69×10^−7^ and LogFC = 1.2383, P = 2.69×10^−7^, FDR = 1.80×10^−4^) respectively. *Adipor2* (Adiponectin receptor 2) expression was upregulated in cluster 1 (LogFC = 0.9853, P = 7.52×10^−7^, FDR = 1.04×10^−3^) (**Table S3**). In the neuronal clusters, pathway analysis revealed that the synaptogenesis signalling pathway is downregulated in cluster 0 in the fasted state (**Table S4**).

#### CALCR

We identified 185 *Calcr* expressing nuclei that were located in 23 of the 30 neuronal clusters (**Figure 5a**). When re-clustered the CALCR neurons grouped into 3 clusters (**Figure 5b**). Cluster 0 and some of cluster 2 expressed *Chrm3* (Acetylcholine muscarinic receptor 3), suggesting that they respond to acetylcholine signalling (**Figure 5c**). Cluster 2 expressed *Th* (tyrosine hydroxylase), suggesting that these are catecholamine neurons (**Figure 5c**).

**Figure 5.**
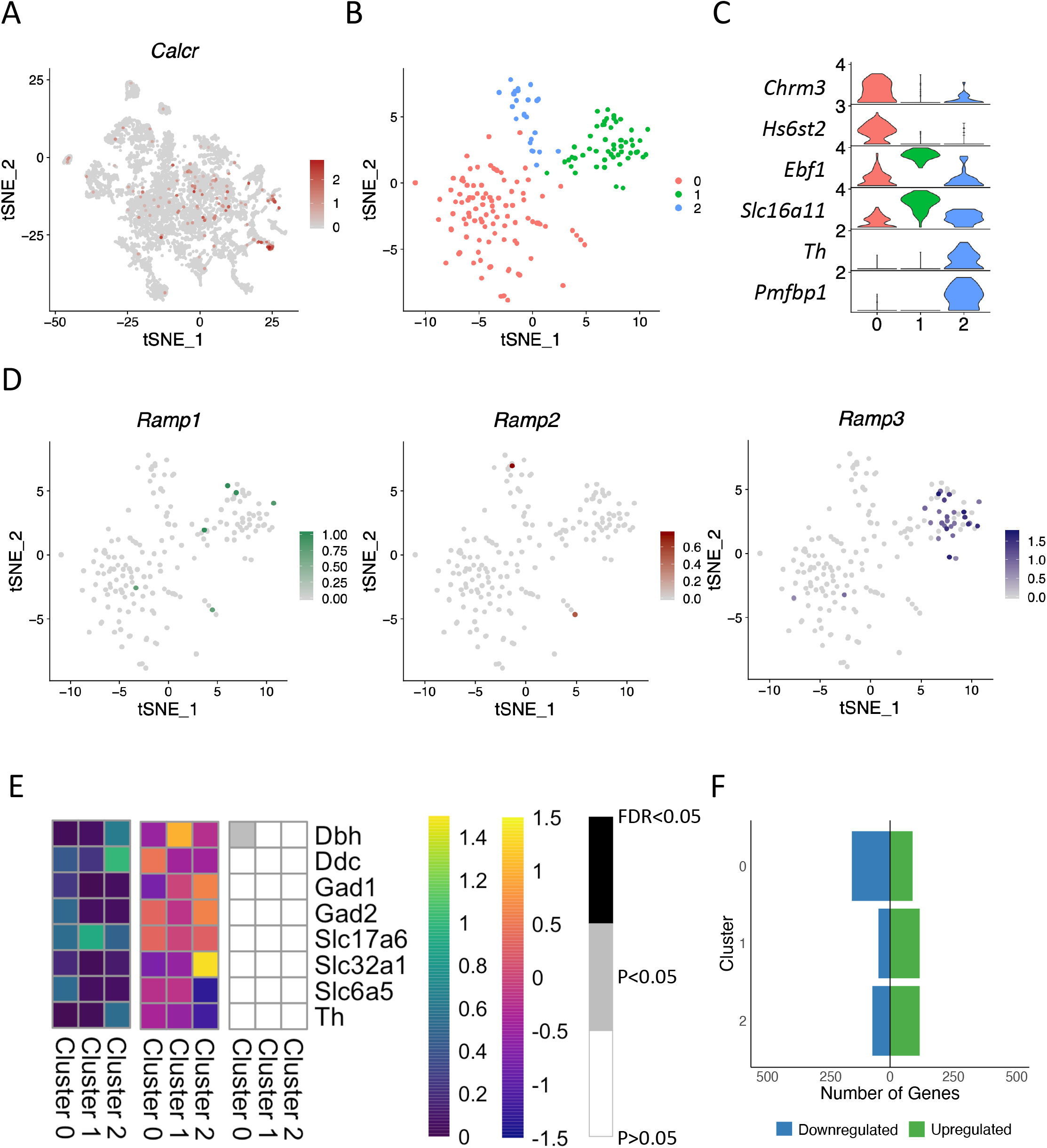
Profiling CALCR neurons in the AP and NTS. (A) Feature plot revealing relative expression levels of *Calcr* in the neuronal clusters. (B) tSNE plot of the 185 CALCR neurons, grouped into 3 clusters. (C) Violin plots showing relative expression levels of 2 marker genes for each cluster of the CALCR neurons. (D) Feature plots showing the relative expression levels of *Ramp1, Ramp2* and *Ramp3* in the CALCR subset. (E) Heatmaps profiling the expression of canonical neuronal markers in each CALCR cluster. Left: Average relative expression level in each cluster; Middle: Differential expression of each gene per cluster (Log2FC values); Right: Determining whether gene expression was significantly changed in the fasted state. (F) Graph showing the number of genes upregulated (green) and downregulated (blue) in response to an overnight fast in each cluster. Genes included were significantly differentially regulated (P<0.05).

Amylin receptors are formed by the heterodimerisation of CALCR with at least one of the 3 Ramp proteins [26]. To identify which of these CALCR neurons are likely expressing the amylin receptor, expression of Ramp1, Ramp2 and Ramp3 was characterised in CALCR neurons (**Figure 5d**). *Ramp1* and *Ramp2* were not widely expressed in the CALCR population, but *Ramp3* was expressed in cluster 1 (**Figure 5d**).

Upon looking at expression of established neuronal markers in each cluster, we see that *Calcr* clusters consist of mainly glutamatergic neurons (**Figure 5e**). Cluster 0 was a mix of both excitatory (43.9%) and inhibitory neurons (31.9%). The expression of *Slc17a6* in cluster 1 suggest that neurons in this cluster were glutamatergic. Cluster 2 nuclei expressed *Slc17a6, Th, Dbh* and *Ddc*, suggesting that this cluster are catecholamine neurons (**Figure 5e**). Detailed expression profiles of these clusters can be found in **Figure S4.**

Cluster 0 exhibited the highest number of differential gene expression in response to a fast (240 genes, **Figure 5f**). In both cluster 0 and cluster 2, *Meg3* expression was downregulated in response to a fast (LogFC = −0.2462, P=8.13×10^−15^; FDR = 2.76×10^−11^ and LogFC = −0.3476, P = 2.23×10^−7^, FDR = 3.77×10^−4^ respectively). Scaffold protein *Kidins220* was downregulated in cluster 2 in the fasted state (LogFC = −2.4617, P = 3.35×10^−9^, FDR = 1.13×10^−5^; **Table S5**).

#### GFRAL

The hormone GDF15 acts to suppress food intake through the GFRAL receptor, located on a small population of neurons within the AP and NTS [27–30]. We identified 114 *Gfral* expressing nuclei. Cluster 31 contained the highest number, with 20.3% of nuclei expressing *Gfral* (**Figure 6a**). A large proportion of GFRAL neurons and GLP1R neurons are located within the same cluster in the overall dataset (cluster 31), indicating the transcriptional similarities between these two populations. However, with our approach we see minimal co-expression of these two receptors, although it is possible that we are not detecting cells expressing low levels of these transcripts.

**Figure 6.**
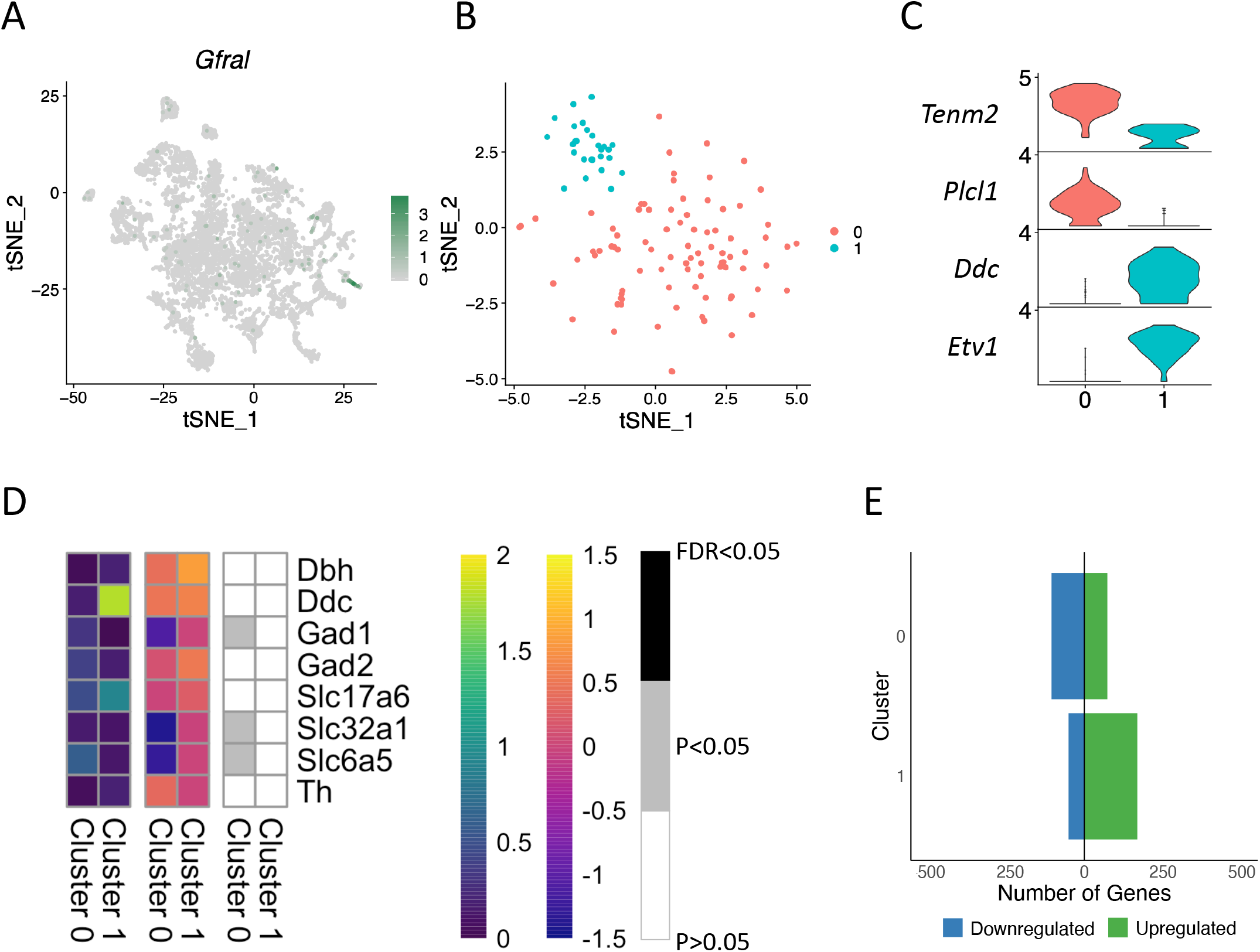
Characterising GFRAL neurons in the AP and NTS. (A) Feature plot of *Gfral* expression in the neuronal dataset. (B) tSNE plot of the 114 GFRAL neurons, grouped into 2 clusters. (C) Violin plots showing relative expression levels of marker genes for both GFRAL clusters. (D) Heatmaps profiling the expression of canonical neuronal markers in both GFRAL clusters. Left: Average relative expression level in each cluster; Middle: Differential expression of each gene per cluster (Log2FC values); Right: Determining whether gene expression was significantly changed in the fasted state. (E) Graph showing the number of genes upregulated (green) and downregulated (blue) in response to an overnight fast in both clusters. Genes included were significantly differentially regulated (P<0.05).

When re-clustered, they form 2 clusters (**Figure 6b**). Cluster 0 was characterised by expression of *Tenm2* and *Plcl1*, while cluster 1 expressed *Ddc*, suggesting that these are dopaminergic neurons (**Figure 6c**). Cluster 1 also expressed *Slc17a6*, *Dbh* and *Th*, suggesting that these neurons transmit both glutamatergic and catecholamine signals (**Figure 6d**).

57 of the 114 *Gfral* neurons came from fasted animals. 177 genes were differentially expressed in the fasted condition in cluster 0, including the inhibitory neuronal markers (*Gad1, Slc32a1* and *Slc6a5*), which were downregulated in the fasted state (LogFC = −1.093, P = 0.0454, FDR = 0.831; LogFC = −1.399, P = 0.00563, FDR = 0.575; and LogFC = −1.282, P = 0.0259, FDR = 0.750 respectively). 118 differentially expressed genes were identified in cluster 1 (**Figure 6e; Table S7**), including *Tenm3* and *Pias1,* that were both upregulated in the fasted condition (LogFC = 2.2753, P = 1.37×10^−5^, FDR = 0.0372 and LogFC = 2.4759, P = 3.33×10^−5^, P = 0.0450 respectively; **Table S7**).

### Characterization of AP/NTS POMC cells

While POMC neurons located within the hypothalamus have been well characterised using single cell RNA-sequencing [16], the NTS POMC neurons have not been similarly studied. Here, we have identified 346 nuclei expressing POMC (**Figure 7**). Unlike in the hypothalamus, where expression of *Pomc* itself appears to be a main driver of cluster formation [14, 15], there was no single cluster that expressed particularly high levels of *Pomc* (**Figure 7a**). Instead, its expression was detected in all clusters apart from the microglia and epithelial cell clusters, with neuronal clusters 40 (*Slc6a4* and *Tph2)*, 36 (*Prph, Tbx20, Chat* and *Slc5a7)* and 22 (*Gal* and *Cartpt*) containing the highest percentage of *Pomc* nuclei (8.7%, 7.2% and 6.1% respectively). Interestingly, *Pomc* expression is also evident throughout the oligodendrocyte clusters. When analysed separately, the 346 nuclei group into 3 neuronal and 1 oligodendrocyte clusters (**Figure 7b**). Clusters 0-2 are neuronal, although their expression profiles are very similar (**Figure 7c**), with cluster 0 nuclei marked by expression of *Baiap3* and *Camk2a*, and clusters 1 and 2 expressing *Syt2* (synaptotagmin) and *Flt3.* Cluster 3 is clearly an oligodendrocyte cluster due to high expression of *Mog*.

**Figure 7.**
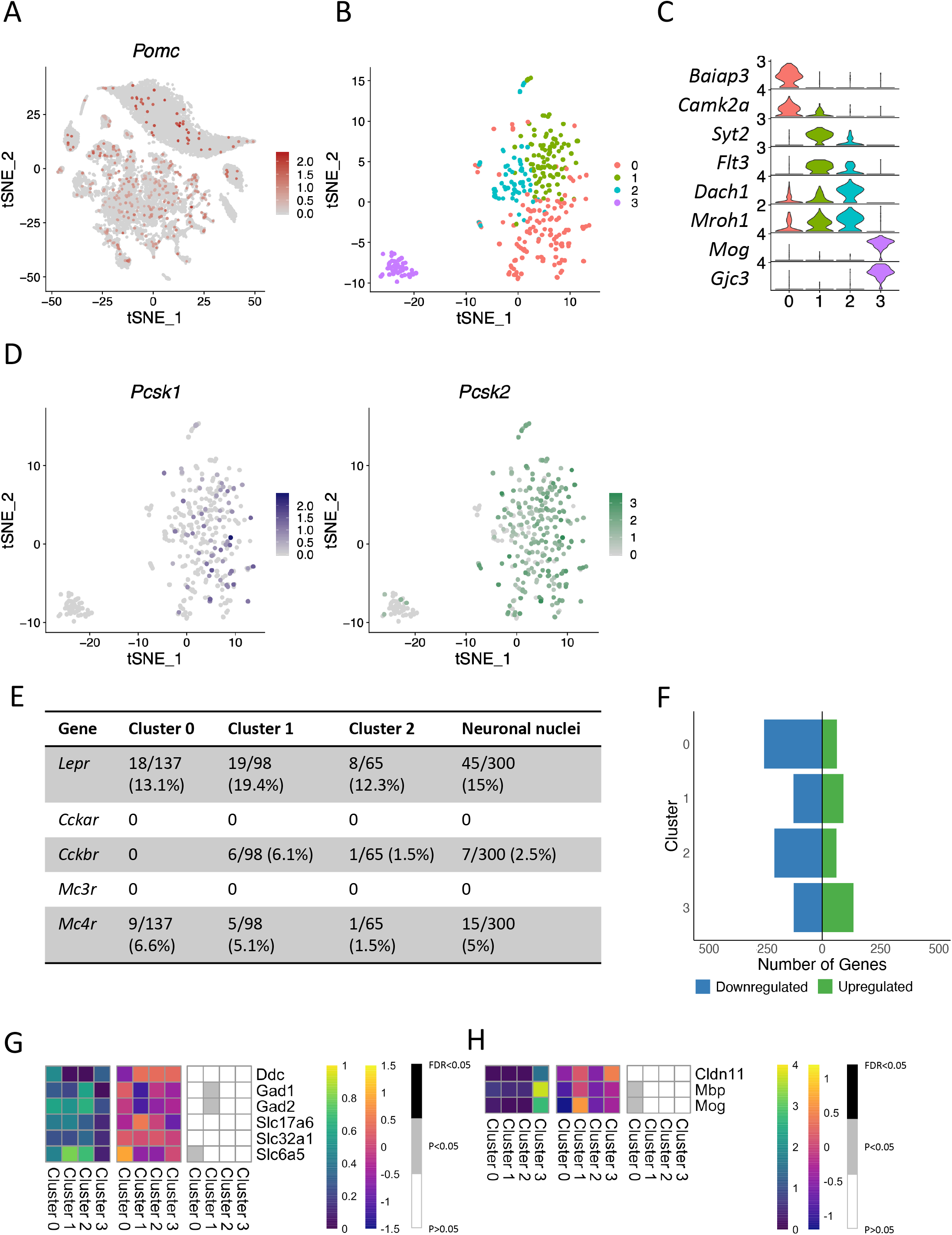
Characterising POMC cells in the AP and NTS. (A) Feature plot showing relative expression levels of *Pomc* in the overall dataset. (B) tSNE plot of the 346 POMC nuclei, grouped into 4 clusters. (C) Violin plots showing relative expression levels of marker genes for each cluster of the POMC subset. (D) The relative expression levels of *Pcsk1* and *Pcsk1* in the POMC subset. (E) Table displaying the number (and percentage) of Pomc neurons expressing *Lepr, Cckar, Cckbr, Mc3r* and *Mc4r* in each of the neuronal clusters, and total expression numbers in the neuronal clusters. (F) Graph showing the number of genes upregulated (green) and downregulated (blue) in response to an overnight fast in each cluster. Genes included were significantly differentially regulated (P<0.05). (G-H) Heatmaps profiling the expression of canonical neuronal (G) and oligodendrocyte (H) markers in each POMC cluster. Left: Average relative expression level in each cluster; Middle: Differential expression of each gene per cluster (Log2FC values); Right: Determining whether gene expression was significantly changed in the fasted state.

POMC is processed into its active constituents by prohormone convertases PCSK1 and PCSK2 [31], and a total of 84% of POMC neuronal nuclei expressed either or both enzymes (**Figure 7d**). In contrast, only four oligodendrocyte nuclei expressed *Pcsk2,* with none expressing *Pcsk1*. Only a small proportion of each cluster expressed the leptin receptor *Lepr* (Cluster 0 = 13.1%, cluster 1 = 19.4% and cluster 2 = 12.3%; **Figure 7e**). 2.5% express the Cck receptor (*Cckbr*) and 5% express *Mc4r*. There was no expression of *Cckar* and *Mc3r* in the POMC nuclei.

In the hypothalamus, POMC neurons are very responsive to changes in nutritional state [14]. In the hindbrain, Cluster 0 exhibited the largest number of differentially expressed genes with the majority being downregulated in the fasted state (**Figure 7f**). This included *Ryr2* (LogFC = −0.7386, P = 6.51×10^−6^, FDR = 0.00569; **Table S9**). 266 genes were differentially regulated in cluster 2. In the oligodendrocyte cluster *Adipor2* expression was upregulated (LogFC = 1.4047, P = 7.44×10^−6^, FDR = 0.00867), and *Frmd4a* was downregulated (LogFC = −3.1281, P = 4.96×10^−14^, FDR = 1.73×10^−10^). In clusters 0 and 2, the synaptogenesis signalling pathway was the top downregulated pathway (**Table S10**).

## Discussion

To date, cells within the AP/NTS have largely been characterised using histological and molecular approaches. In this study, we have performed a single-nucleus level survey of 16,034 cells mostly from the mouse AP and NTS in the fed and fasted state. Because of our method of dissection, we undoubtedly have also captured cells from adjacent nuclei such as the dorsal motor nucleus of the vagus. Here we provide detailed expression profiles of hindbrain cells expressing the key anti-obesity drug targets GLP1R, GIPR, CALCR and GFRAL, and were able to capture and profile NTS POMC neurons at a single nucleus level.

A key benefit of NucSeq is that it can be performed on CNS tissue from older animals, as neurons in older animals tend to have longer processes and are likely to be damaged in the single cell dissociation process [32]. At best, this can result in sampling biases due to cells with long processes being underrepresented in the dataset, while at worst nothing is captured at all. Thus, NucSeq opens up the possibility of studying the effects of long-term high fat diet or other diets on changes in gene expression. Furthermore, as a cell nucleus turns out to be a pretty robust organelle, NucSeq can be performed on frozen samples, thereby substantially expanding the availability of suitable samples to include those archived in tissue banks. This will certainly be the preferred method to use on human brain samples [33], and thus NucSeq data from mouse samples would be a more appropriate comparator for human single brain cell transcriptomics. While cytosolic RNA is inherently different from the mix of unspliced RNA emerging from the transcriptional machinery within the nucleus, transcriptome data from single nuclei are qualitatively reflective of that from single cells, certainly in terms of different cell types and their unique repertoire of transcripts [34].

Previous transcriptomic analyses of the mouse brain in the fed and fasted state have focussed on the hypothalamus, either as a whole, on individually laser capture microdissected regions, or most recently using single cell/nucleus RNA-sequencing [15, 35]. From these studies, we know that there are many different populations of hypothalamic neurons that are exquisitely sensitive to changes in nutritional state [14, 15]. In contrast, here, we show that hindbrain neuronal populations were not especially responsive to an overnight fast, with the majority exhibiting modest effects on differential gene expression. This is not a surprise, and reflects the differences in physiological roles of these two regions in the brain when it comes to appetitive drive, with the hypothalamus regulating hunger and satiety, and the hindbrain mediating the more visceral sensations, ranging from satiety and fullness, to aversion and nausea [3]. Unexpectedly, the one exception was the population of hindbrain oligodendrocytes, that we show to be transcriptionally sensitive to the fasted state. In particular, using pathway analysis, we found that an upstream regulator of differentially expressed genes in oligodendrocytes was the transcription factor *Tcf7l2*, which has been previously shown to be necessary for differentiation of oligodendrocytes [36]. This suggests a possible remodelling of hindbrain oligodendrocytes that occurs in response to an overnight fast, a phenomenon that has previously been seen in oligodendrocytes in the median eminence (ME) of the hypothalamus [37]. Like in the ME, the hindbrain oligodendrocytes are positioned near a circumventricular organ, and we suggest they could be, in response to nutritional signals, controlling access of circulating metabolic cues to neurons in the AP.

The role of the hindbrain in mediating the satiety and aversive responses is part of the reason why many anti-obesity therapies are effective at the hindbrain. For example, once weekly semaglutide administration over a period of 15 months has recently been shown to produce a mean bodyweight reduction of 14.9% in patients with obesity [5]. Additional to signalling to hypothalamic neurons, semaglutide and liraglutide have both been shown to activate GLP1R neurons within the AP and NTS, mediating some of the weight loss effects of these drugs [7, 38]. The four neuronal populations we focus on, expressing GLP1R, GIPR, CALCR and GFRAL, appear to be distinct of each other, possibly indicating that each of these receptors signals through independent pathways. However, it is important to note that while NucSeq is effective at identifying the presence of transcripts it is not designed to demonstrate the absence of a transcript. Thus, it is entirely possible that there are cells co-expressing these receptors at low levels that we are not detecting.

Even with dual agonism of GIPR and GLP1R showing an additional effect on weight loss in humans by reduction of food intake [39], the involvement of GIPR in the control of food intake and bodyweight is still not fully understood, with both antagonism of the receptor with an antibody, and administration, either centrally or peripherally, of a GIPR agonist showing efficacious results in weight loss in mice [40, 41]. While it is clear that hypothalamic GIPR plays a key role in this effect on weight loss; what is less clear is the role of GIPR in the hindbrain. For one thing, unlike GLP1R, whose expression in the hindbrain was largely restricted to neurons, around half of the cells expressing GIPR are oligodendrocytes. Like the broader population of oligodendrocytes, the subset of GIPR expressing cells are also transcriptionally sensitive to an overnight fast, although it is, as yet, unclear the role GIPR signalling plays in these cells.

Another class of compounds that work in concert with GLP1R agonists are amylin mimetics, with dual amylin and calcitonin receptor agonists shown to have complimentary actions on food intake in combination with liraglutide treatment [42]. CALCR neurons within the NTS are a key mediator of food intake reduction from CALCR agonists in comparison with the hypothalamus [43], demonstrating the importance of characterising these hindbrain populations. Of the three distinct populations of CALCR neurons we identify, only one is likely to express the amylin receptor based on its co-expression of *Ramp3*.

Finally, with approval of the melanocortin agonist setmelanotide for treatment of rare causes of obesity, there has been renewed interest in the hypothalamic melanocortin pathway. But the POMC neurons in the hindbrain, while they have been studied neurochemically, have never been successfully isolated and characterised using transcriptomic techniques. Previously, POMC mRNA expression in the NTS has not been well characterised due to expression levels being so low, consequently it has been difficult to detect at a single cell level [44]. Here we have been able to identify and characterise POMC neurons within the hindbrain and show that the majority of these neurons co-express pro-hormone convertases PCSK1 and/or PCSK2. A number of functional studies have demonstrated that leptin administration activates POMC neurons in the NTS through elevation of c-fos or pSTAT3 immunoreactivity [44, 45] pointing to the role of these neurons in the regulation of satiety signalling. Indeed, there are contradictory findings on the expression of LepRb on POMC NTS neurons, with studies using POMC-eGFP mice demonstrating that a subpopulation of POMC neurons express LepRb, and a study using a POMC-Cre reporter mouse finding no co-expression of POMC with LepRb [44–46]. In our dataset, we find that a small subpopulation of POMC neurons (15%) do in fact express *Lepr* transcripts. A possible explanation for some of the inconsistent findings regarding the properties of POMC neurons within the NTS are the differences in which cells are marked by POMC-eGFP and POMC-Cre mouse lines [47]. Identification of POMC neurons through RNA-sequencing will help to clarify and strengthen evidence surrounding the expression profiles of these neurons.

To conclude, we report here single-nucleus RNA-sequencing of 30 neuronal, and 11 non neuronal populations residing within the AP/NTS. In addition, we provide extensive profiling of differential gene expression that occurs in response to an overnight fast and identify that oligodendrocytes in fact exhibit the greatest changes in transcriptional expression. We have provided detailed profiling of GLP-1R, CALCR, GIPR and GFRAL cells, based on their expression of receptors, ion channels and transporters as well as their neuronal/glial properties, informing future studies that will study these particular targets. Finally, for the first time we have been able to identify and characterise POMC cells residing within the AP/NTS on the single cell level. This resource will help delineate the mechanisms underlying the effectiveness of these compounds, and also prove useful in the continued search for other novel therapeutic targets for obesity.

## Author Contributions

G.K.C.D collected, analysed and visualized data, wrote original draft and edited manuscript; B.Y.H.L collected, analysed and visualized data, and edited manuscript; J.T., I.C., D.R. collected data and edited manuscript; A.P.C. supervision and edited manuscript; J.P.W., L.B.K. data analysis and edited manuscript; C.P. funding acquisition, supervision and edited manuscript; G.S.H.Y conceptualisation, funding acquisition, supervision, wrote original draft, reviewed and edited manuscript.

---

**Supplementary Figure 1**

Characterisation of neuronal populations from the AP and NTS. A heatmap of neuronal cluster marker transcript expression. Top markers for each cluster were determined using Wilcoxon Ranked Sum Test and ROC analysis. Cluster numbers are displayed across the top of the heatmap. Colours representing each cluster are consistent with figure 1D.

**Supplementary Figure 2**

Expression profiles of GLP1R neurons. Heatmaps showing the average relative expression levels (left), differential gene expression (centre), and significance values (right) of genes encoding: GPCRs, nucleus transporters, plasma membrane transporters, cytoplasm transporters, ion channels, kinase receptors, transmembrane receptors and peptides. Genes were included if transcripts were expressed in at least 50% of nuclei in one cluster. Left panels: average relative expression levels of each gene in each cluster, with low/no expression in deep blues and high expression in greens. Middle: Log fold change of gene expression in each cluster in response to an overnight fast. No change in gene expression in pinks, downregulated gene expression in purples, upregulated expression in yellow/orange. Right: Determining the significance of differential gene expression in response to an overnight fast in each cluster. White: P>0.05; grey: P<0.05; black: FDR < 0.05.

**Supplementary Figure 3**

Characterisation of GIPR nuclei. Heatmaps showing the average relative expression levels (left), differential gene expression (centre), and significance values (right) of genes encoding: GPCRs, nucleus transporters, plasma membrane transporters, cytoplasm transporters, ion channels, kinase receptors, transmembrane receptors and peptides. Left panels: average relative expression levels of each gene in each cluster, with low/no expression in deep blues and high expression in greens. Middle: Log fold change of gene expression in each cluster in response to an overnight fast. No change in gene expression in pinks, downregulated gene expression in purples, upregulated expression in yellow/orange. Right: Determining the significance of differential gene expression in response to an overnight fast in each cluster. White: P>0.05; grey: P<0.05; black: FDR < 0.05.

**Supplementary Figure 4**

Characterisation of CALCR neurons. Heatmaps showing the average relative expression levels (left), differential gene expression (centre), and significance values (right) of genes encoding: GPCRs, nucleus transporters, plasma membrane transporters, cytoplasm transporters, ion channels, kinase receptors, transmembrane receptors and peptides. Left panels: average relative expression levels of each gene in each cluster, with low/no expression in deep blues and high expression in greens. Middle: Log fold change of gene expression in each cluster in response to an overnight fast. No change in gene expression in pinks, downregulated gene expression in purples, upregulated expression in yellow/orange. Right: Determining the significance of differential gene expression in response to an overnight fast in each cluster. White: P>0.05; grey: P<0.05; black: FDR < 0.05.

**Supplementary Figure 5**

Characterisation of GFRAL neurons. Heatmaps showing the average relative expression levels (left), differential gene expression (centre), and significance values (right) of genes encoding: GPCRs, nucleus transporters, plasma membrane transporters, cytoplasm transporters, ion channels, kinase receptors, transmembrane receptors and peptides. Left panels: average relative expression levels of each gene in each cluster, with low/no expression in deep blues and high expression in greens. Middle: Log fold change of gene expression in each cluster in response to an overnight fast. No change in gene expression in pinks, downregulated gene expression in purples, upregulated expression in yellow/orange. Right: Determining the significance of differential gene expression in response to an overnight fast in each cluster. White: P>0.05; grey: P<0.05; black: FDR < 0.05.

**Supplementary Figure 6**

Profiling POMC nuclei. Heatmaps showing the average relative expression levels (left), differential gene expression (centre), and significance values (right) of genes encoding: GPCRs, nucleus transporters, plasma membrane transporters, cytoplasm transporters, ion channels, kinase receptors, transmembrane receptors and peptides. Left panels: average relative expression levels of each gene in each cluster, with low/no expression in deep blues and high expression in greens. Middle: Log fold change of gene expression in each cluster in response to an overnight fast. No change in gene expression in pinks, downregulated gene expression in purples, upregulated expression in yellow/orange. Right: Determining the significance of differential gene expression in response to an overnight fast in each cluster. White: P>0.05; grey: P<0.05; black: FDR < 0.05.

**Supplementary Table 1**

Top differentially regulated genes in each GLP1R cluster in response to an overnight fast. Genes included in analysis were expressed in at least 50% of one cluster.

**Supplementary Table 2**

Top 5 canonical pathways affected by an overnight fast in GLP1R clusters. Data taken from Ingenuity Pathway analysis. Overlap denotes the number of genes in that pathway that were differentially regulated.

**Supplementary Table 3**

Top differentially regulated genes in each GIPR cluster in response to an overnight fast.

**Supplementary Table 4**

Top 5 canonical pathways affected in each GIPR cluster by an overnight fast. Overlap denotes the number of genes in that pathway that were differentially regulated.

**Supplementary Table 5**

Top differentially regulated genes in each CALCR cluster in response to an overnight fast.

**Supplementary Table 6**

Top 5 canonical pathways affected in CALCR clusters by an overnight fast. Overlap denotes the number of genes in that pathway that were differentially regulated.

**Supplementary Table 7**

Top differentially regulated genes in both GFRAL clusters in response to an overnight fast.

**Supplementary Table 8**

Top 5 canonical pathways affected in both clusters by an overnight fast in the GFRAL subset. Overlap denotes the number of genes in that pathway that were differentially regulated.

**Supplementary Table 9**

Top differentially regulated genes in each POMC cluster in response to an overnight fast.

**Supplementary Table 10**

Top 5 canonical pathways affected in each cluster by an overnight fast in the POMC subset. Overlap denotes the number of genes in that pathway that were differentially regulated.

