## Supplemental tables for "A survey of the mouse hindbrain in the fed and fasted state using single-nucleus RNA sequencing"

### Supplementary Table 1

| Cluster | Gene | LogFC | Pvalue | FDR | Upregulated or downregulated |
|---|---|---|---|---|---|
| 0 | *Ptgds* | 1.1566 | 6.40E-08 | 0.0003 | Upregulated |
| 0 | *Srsf10* | 0.936 | 1.19E-05 | 0.0099 | Upregulated |
| 0 | *Hs3st2* | 2.0424 | 1.45E-04 | 0.0749 | Upregulated |
| 0 | *Zmym1* | -1.3848 | 7.38E-07 | 0.0015 | Downregulated |
| 0 | *CT030170.6* | -0.9617 | 4.09E-06 | 0.0048 | Downregulated |
| 0 | *Nudcd3* | -0.8974 | 4.62E-06 | 0.0048 | Downregulated |
| 0 | *Col26a1* | -1.3395 | 8.35E-05 | 0.0576 | Downregulated |
| 0 | *Ring1* | -0.8939 | 1.09E-04 | 0.0644 | Downregulated |
| 0 | *Igf2bp3* | -0.7801 | 2.79E-04 | 0.1163 | Downregulated |
| 0 | *Tsga10* | -1.1958 | 2.81E-04 | 0.1163 | Downregulated |
| 1 | *Fgf1* | 2.9815 | 2.76E-04 | 0.2286 | Upregulated |
| 1 | *Clcn4* | 2.3135 | 3.97E-04 | 0.2714 | Upregulated |
| 1 | *Itch* | 2.3085 | 5.01E-04 | 0.2714 | Upregulated |
| 1 | *Oxr1* | -1.9133 | 3.78E-06 | 0.0156 | Downregulated |
| 1 | *Kcnj3* | -1.9911 | 9.29E-05 | 0.1465 | Downregulated |
| 1 | *Tmeff2* | -1.9994 | 1.06E-04 | 0.1465 | Downregulated |
| 1 | *Vmp1* | -2.2196 | 1.85E-04 | 0.1913 | Downregulated |
| 1 | *1700024B18Rik* | -2.4707 | 6.24E-04 | 0.2714 | Downregulated |
| 1 | *Cdh20* | -2.878 | 6.27E-04 | 0.2714 | Downregulated |
| 1 | *Tpi1* | -2.6634 | 8.27E-04 | 0.2714 | Downregulated |
| 2 | *Trpc5* | 3.837 | 3.01E-04 | 0.3903 | Upregulated |
| 2 | *Dnajc13* | 2.3646 | 3.89E-04 | 0.3903 | Upregulated |
| 2 | *Cops3* | 3.028 | 5.66E-04 | 0.3903 | Upregulated |
| 2 | *Arhgap26* | 2.8088 | 8.31E-04 | 0.4671 | Upregulated |
| 2 | *Csmd1* | 1.6243 | 9.03E-04 | 0.4671 | Upregulated |
| 2 | *Snrnp48* | -2.0124 | 1.04E-04 | 0.3903 | Downregulated |
| 2 | *Gls* | -1.1049 | 5.23E-04 | 0.3903 | Downregulated |
| 2 | *Srsf5* | -2.2289 | 5.57E-04 | 0.3903 | Downregulated |
| 2 | *Cabin1* | -1.6284 | 1.21E-03 | 0.5158 | Downregulated |
| 2 | *Hsp90ab1* | -1.8762 | 1.25E-03 | 0.5158 | Downregulated |

### Supplementary Table 2

| Cluster | Pathway name | Overlap | P value | Z-score |
|---|---|---|---|---|
| 0 | FcyRIIB Signaling in B Lymphocytes | 6/75 | 4.87E-04 | -0.447 |
| 0 | Fc Epsilon RI Signaling | 7/117 | 9.71E-04 | -0.378 |
| 0 | Synaptogenesis Signaling Pathway | 12/312 | 1.02E-03 | -0.302 |
| 0 | Reelin Signaling in Neurons | 7/122 | 1.24E-03 | 1.89 |
| 0 | Regulation of IL-2 Expression in Activated and Anergic T Lymphocytes | 6/90 | 1.28E-03 | N/A |
| 1 | HIPPO Signaling | 6/85 | 1.12E-03 | 0.000 |
| 1 | Protein Ubiquination Pathways | 11/273 | 1.46E-03 | N/A |
| 1 | Ephrin A Signaling | 4/47 | 3.90E-03 | N/A |
| 1 | IL-15 Production | 6/121 | 6.54E-03 | 0.000 |
| 1 | p70S6K Signaling | 6/129 | 8.85E-03 | -0.816 |
| 2 | Glutamate Receptor Signaling | 5/57 | 6.09E-05 | N/A |
| 2 | Calcium Signaling | 8/206 | 1.43E-04 | 1.633 |
| 2 | nNOS Signaling in Skeletal Muscle Cells | 4/41 | 2.27E-04 | N/A |
| 2 | Opioid Signaling Pathway | 8/247 | 4.86E-04 | 2.121 |
| 2 | Synaptogenesis Signaling Pathway | 7/312 | 8.26E-03 | 0.816 |

N/A = Not available

### Supplementary Table 3

| Cluster | Gene | LogFC | Pvalue | FDR | Upregulated or downregulated |
|---|---|---|---|---|---|
| 0 | *Ptgds* | 1.4381 | 3.14E-10 | 8.69E-07 | Upregulated |
| 0 | *Srsf10* | 0.7647 | 6.79E-05 | 6.26E-02 | Upregulated |
| 0 | *Haghl* | -0.8584 | 2.47E-05 | 3.42E-02 | Downregulated |
| 0 | *A230103L15Rik* | -0.6843 | 1.20E-04 | 7.24E-02 | Downregulated |
| 0 | *Brd9* | -0.8472 | 1.54E-04 | 7.24E-02 | Downregulated |
| 0 | *Sox2ot* | -1.1124 | 1.70E-04 | 7.24E-02 | Downregulated |
| 0 | *Kidins220* | -0.6125 | 1.83E-04 | 7.24E-02 | Downregulated |
| 0 | *Thrb* | -0.8053 | 2.26E-04 | 7.74E-02 | Downregulated |
| 0 | *Washc4* | -0.8536 | 2.76E-04 | 7.74E-02 | Downregulated |
| 0 | *Cspp1* | -0.5853 | 2.80E-04 | 7.74E-02 | Downregulated |
| 1 | *Garnl3* | 1.7049 | 2.74E-10 | 7.57E-07 | Upregulated |
| 1 | *Adipor2* | 0.9853 | 7.52E-07 | 1.04E-03 | Upregulated |
| 1 | *Sgk1* | 1.6912 | 1.42E-05 | 6.71E-03 | Upregulated |
| 1 | *Sgk3* | 1.2952 | 1.53E-05 | 6.71E-03 | Upregulated |
| 1 | *Fam13c* | 1.4127 | 1.70E-05 | 6.71E-03 | Upregulated |
| 1 | *A330015K06Rik* | 1.0733 | 3.49E-05 | 1.12E-02 | Upregulated |
| 1 | *AI314180* | 1.2144 | 3.63E-05 | 1.12E-02 | Upregulated |
| 1 | *Fam214a* | 1.4617 | 5.72E-05 | 1.58E-02 | Upregulated |
| 1 | *Ncam2* | -0.9466 | 1.13E-05 | 6.71E-03 | Downregulated |
| 1 | *Prkca* | -1.2076 | 1.62E-05 | 6.71E-03 | Downregulated |
| 2 | *Ptgds* | 1.2383 | 2.69E-07 | 1.80E-04 | Upregulated |
| 2 | *Map7d2* | 1.9026 | 8.52E-06 | 2.36E-03 | Upregulated |
| 2 | *Frmd4a* | -3.5686 | 1.96E-27 | 5.43E-24 | Downregulated |
| 2 | *Lsamp* | -1.7686 | 3.42E-13 | 4.72E-10 | Downregulated |
| 2 | *Tnr* | -2.474 | 7.33E-08 | 6.76E-05 | Downregulated |
| 2 | *Tmem108* | -1.8941 | 3.26E-07 | 1.80E-04 | Downregulated |
| 2 | *Ptprj* | -2.2109 | 4.18E-07 | 1.92E-04 | Downregulated |
| 2 | *Pdcd4* | -1.9906 | 1.28E-06 | 5.06E-04 | Downregulated |
| 2 | *Opcml* | -1.564 | 2.68E-06 | 9.27E-04 | Downregulated |
| 2 | *Cadm2* | -1.0895 | 7.05E-06 | 2.16E-03 | Downregulated |
| 3 | *Mapre3* | 2.5904 | 5.46E-04 | 3.77E-01 | Upregulated |
| 3 | *Papola* | 1.9635 | 7.18E-04 | 3.77E-01 | Upregulated |
| 3 | *Atad1* | 2.5059 | 8.18E-04 | 3.77E-01 | Upregulated |
| 3 | *Ranbp2* | 2.1467 | 1.75E-03 | 5.21E-01 | Upregulated |
| 3 | *Mgat4c* | 2.2483 | 1.78E-03 | 5.21E-01 | Upregulated |
| 3 | *Rnf130* | -1.6106 | 1.69E-04 | 3.77E-01 | Downregulated |
| 3 | *Gpc6* | -2.2961 | 3.82E-04 | 3.77E-01 | Downregulated |
| 3 | *Scfd1* | -1.7328 | 7.56E-04 | 3.77E-01 | Downregulated |
| 3 | *Phf21a* | -1.225 | 1.96E-03 | 5.21E-01 | Downregulated |
| 3 | *BC052040* | -1.8271 | 2.61E-03 | 5.21E-01 | Downregulated |

### Supplementary Table 4

| Cluster | Pathway name | Overlap | P value | Z-score |
|---|---|---|---|---|
| 0 | Insulin Secretion Signalling Pathway | 10/244 | 9.31E-05 | -1.265 |
| 0 | Glutamate Receptor Signaling | 4/57 | 1.88E-03 | N/A |
| 0 | Synaptic Long Term Depression | 6/189 | 8.23E-03 | -0.816 |
| 0 | Synaptogenesis Signaling Pathway | 8/312 | 8.52E-03 | -1.414 |
| 0 | Iron Homeostasis signaling pathway | 5/137 | 8.83E-03 | N/A |
| 1 | Melatonin Signaling | 7/72 | 1.94E-05 | -0.378 |
| 1 | Role of Tissue Factor in Cancer | 8/116 | 6.09E-05 | N/A |
| 1 | Relaxin Signaling | 9/151 | 6.66E-05 | N/A |
| 1 | Synaptic Long Term Depression | 10/189 | 7.21E-05 | 0.333 |
| 1 | G-Protein Coupled Receptor Signaling | 12/274 | 8.71E-05 | N/A |
| 2 | Reelin Signaling in Neurons | 8/122 | 1.87E-04 | -0.707 |
| 2 | Axonal Guidance Signaling | 17/494 | 2.52E-04 | N/A |
| 2 | Semaphorin Neuronal Repulsive Signaling Pathway | 8/139 | 4.53E-04 | -0.707 |
| 2 | RhoGDI Signaling | 9/189 | 8.04E-04 | 1.414 |
| 2 | Glutamate Receptor Signaling | 5/57 | 8.31E-04 | N/A |
| 3 | Pyridoxal 5’-phosphate Salvage Pathway | 4/66 | 2.00E-03 | 1.000 |
| 3 | Agrin Interactions at Neuromuscular Junction | 4/70 | 2.48E-03 | 1.000 |
| 3 | IL-15 Production | 5/121 | 3.02E-03 | 0.447 |
| 3 | Cleavage and Polyadenylation of Pre-mRNA | 2/12 | 4.05E-03 | N/A |
| 3 | ErbB Signaling | 4/94 | 7.10E-03 | 2.000 |

N/A = Not available

### Supplementary Table 5

| Cluster | Gene | LogFC | Pvalue | FDR | Upregulated or downregulated |
|---|---|---|---|---|---|
| 0 | *Srsf10* | 1.0494 | 1.65E-05 | 2.79E-02 | Upregulated |
| 0 | *Ptbp2* | 0.8413 | 5.43E-04 | 4.60E-01 | Upregulated |
| 0 | *Psmb2* | 0.8812 | 1.11E-03 | 4.77E-01 | Upregulated |
| 0 | *Rps6ka5* | 1.1784 | 2.00E-03 | 4.77E-01 | Upregulated |
| 0 | *Meg3* | -0.2462 | 8.13E-15 | 2.76E-11 | Downregulated |
| 0 | *Erlec1* | -0.9549 | 4.27E-04 | 4.60E-01 | Downregulated |
| 0 | *Ppp2r2c* | -0.97 | 8.18E-04 | 4.77E-01 | Downregulated |
| 0 | *Dach1* | -1.2887 | 1.38E-03 | 4.77E-01 | Downregulated |
| 0 | *Csmd2* | -0.6804 | 1.54E-03 | 4.77E-01 | Downregulated |
| 0 | *Fgf14* | -0.307 | 2.16E-03 | 4.77E-01 | Downregulated |
| 1 | *Ints6* | 1.7561 | 3.34E-04 | 3.16E-01 | Upregulated |
| 1 | *Ptpn4* | 1.4941 | 4.26E-04 | 3.16E-01 | Upregulated |
| 1 | *Ppp3ca* | 1.2751 | 7.31E-04 | 3.16E-01 | Upregulated |
| 1 | *Dnajc10* | 1.812 | 7.34E-04 | 3.16E-01 | Upregulated |
| 1 | *Eif3j1* | 2.1708 | 8.23E-04 | 3.16E-01 | Upregulated |
| 1 | *Ubxn7* | 1.6836 | 9.31E-04 | 3.16E-01 | Upregulated |
| 1 | *Desi1* | 1.6832 | 9.33E-04 | 3.16E-01 | Upregulated |
| 1 | *Osbpl8* | -1.9037 | 1.08E-04 | 3.16E-01 | Downregulated |
| 1 | *Map2k2* | -1.5608 | 6.38E-04 | 3.16E-01 | Downregulated |
| 1 | *BC065397* | -1.8 | 7.93E-04 | 3.16E-01 | Downregulated |
| 2 | *Izumo4* | 1.753 | 9.40E-04 | 4.55E-01 | Upregulated |
| 2 | *Stim1* | 2.2095 | 1.39E-03 | 4.86E-01 | Upregulated |
| 2 | *Kidins220* | -2.4617 | 3.35E-09 | 1.13E-05 | Downregulated |
| 2 | *Meg3* | -0.3476 | 2.23E-07 | 3.77E-04 | Downregulated |
| 2 | *Arhgap26* | -3.2975 | 1.69E-05 | 1.59E-02 | Downregulated |
| 2 | *Cntnap5b* | -2.5832 | 1.88E-05 | 1.59E-02 | Downregulated |
| 2 | *Cntn6* | -2.7569 | 5.77E-05 | 3.91E-02 | Downregulated |
| 2 | *Agfg1* | -2.0325 | 4.57E-04 | 2.58E-01 | Downregulated |
| 2 | *Pcdh17* | -1.5186 | 1.32E-03 | 4.86E-01 | Downregulated |
| 2 | *Pde10a* | -1.1658 | 1.43E-03 | 4.86E-01 | Downregulated |

### Supplementary Table 6

| Cluster | Pathway name | Overlap | P value | Z-score |
|---|---|---|---|---|
| 0 | ILK Signaling | 11/190 | 2.98E-06 | 0.905 |
| 0 | CDK5 Signaling | 6/108 | 6.97E-04 | -0.447 |
| 0 | Synaptogenesis Signaling Pathway | 10/312 | 1.08E-03 | 0.632 |
| 0 | BEX2 Signaling Pathway | 5/79 | 1.09E-03 | 0.447 |
| 0 | TR/RXR Activation | 5/84 | 1.43E-03 | N/A |
| 1 | PTEN Signaling | 8/136 | 3.87E-05 | -1.414 |
| 1 | Cardiac Hypertrophy Signaling (Enhanced) | 15/497 | 6.15E-05 | 2.840 |
| 1 | PI3K Signaling in B Lymphocytes | 7/138 | 2.96E-05 | 1.890 |
| 1 | B Cell Receptor Signaling | 8/186 | 3.39E-04 | 1.414 |
| 1 | Opioid Signaling Pathway | 9/247 | 4.39E-04 | 1.000 |
| 2 | NGF Signaling | 7/114 | 2.76E-05 | -1.134 |
| 2 | Apelin Endothelial Signaling Pathway | 7/115 | 2.91E-05 | -1.134 |
| 2 | Ovarian Cancer Signaling | 7/139 | 9.74E-05 | -0.447 |
| 2 | Estrogen Receptor Signaling | 10/328 | 2.22E-04 | -1.000 |
| 2 | Glucocorticoid Receptor Signaling | 12/426 | 2.34E-04 | N/A |

N/A = Not available

### Supplementary Table 7

| Cluster | Gene | LogFC | Pvalue | FDR | Upregulated or downregulated |
|---|---|---|---|---|---|
| 0 | *Vat1l* | 1.22 | 3.82E-04 | 1.57E-01 | Upregulated |
| 0 | *Rbbp7* | 1.2681 | 7.56E-04 | 2.01E-01 | Upregulated |
| 0 | *Pnisr* | -0.7314 | 1.08E-04 | 1.57E-01 | Downregulated |
| 0 | *Htr1f* | -1.2235 | 1.82E-04 | 1.57E-01 | Downregulated |
| 0 | *Tra2a* | -0.6291 | 2.13E-04 | 1.57E-01 | Downregulated |
| 0 | *Rap1gap* | -0.9628 | 2.74E-04 | 1.57E-01 | Downregulated |
| 0 | *Prpf4b* | -0.4993 | 3.19E-04 | 1.57E-01 | Downregulated |
| 0 | *CT030170.6* | -0.8353 | 4.06E-04 | 1.57E-01 | Downregulated |
| 0 | *Rbm28* | -0.9001 | 8.02E-04 | 2.01E-01 | Downregulated |
| 0 | *Cacnb2* | -0.6172 | 8.71E-04 | 2.01E-01 | Downregulated |
| 1 | *Tenm3* | 2.2753 | 1.37E-05 | 3.72E-02 | Upregulated |
| 1 | *Pias1* | 2.4759 | 3.33E-05 | 4.50E-02 | Upregulated |
| 1 | *Hecw2* | 2.2834 | 4.30E-04 | 2.39E-01 | Upregulated |
| 1 | *4930555F03Rik* | 2.9126 | 4.68E-04 | 2.39E-01 | Upregulated |
| 1 | *Gtdc1* | 2.9365 | 6.38E-04 | 2.47E-01 | Upregulated |
| 1 | *Slc7a14* | 1.9889 | 1.12E-03 | 3.80E-01 | Upregulated |
| 1 | *Gm26691* | 1.9291 | 1.47E-03 | 3.91E-01 | Upregulated |
| 1 | *Pde4b* | -1.6846 | 9.73E-05 | 8.77E-02 | Downregulated |
| 1 | *Wdr26* | -2.9457 | 5.29E-04 | 2.39E-01 | Downregulated |
| 1 | *Srpk1* | -2.4043 | 1.72E-03 | 3.91E-01 | Downregulated |

### Supplementary Table 8

| Cluster | Pathway name | Overlap | P value | Z-score |
|---|---|---|---|---|
| 0 | G-protein Coupled Receptor Signaling | 11/274 | 5.59E-06 | N/A |
| 0 | Axonal Guidance Signaling | 12/494 | 2.78E-04 | N/A |
| 0 | cAMP-mediated signaling | 8/229 | 2.82E-04 | -0.707 |
| 0 | Molecular Mechanisms of Cancer | 10/400 | 7.22E-04 | N/A |
| 0 | T Cell Receptor Signaling | 5/106 | 1.08E-03 | N/A |
| 1 | Synaptogenesis Signaling Pathway | 16/312 | 2.54E-08 | 2.324 |
| 1 | Ephrin Receptor Signaling | 9/189 | 5.56E-05 | 0.816 |
| 1 | Protein Kinase A Signaling | 13/400 | 7.20E-05 | 1.387 |
| 1 | Opioid Signaling Pathway | 9/247 | 4.14E-04 | 1.667 |
| 1 | cAMP-mediated Signaling | 8/229 | 1.13E-03 | 0.707 |

N/A = Not available

### Supplementary Table 9

| Cluster | Gene | LogFC | Pvalue | FDR | Upregulated or downregulated |
|---|---|---|---|---|---|
| 0 | *Ctnna2* | -0.6324 | 1.36E-06 | 3.18E-03 | Downregulated |
| 0 | *Kidins220* | -0.9952 | 2.58E-06 | 3.18E-03 | Downregulated |
| 0 | *Arl15* | -1.0367 | 2.73E-06 | 3.18E-03 | Downregulated |
| 0 | *Ryr2* | -0.7386 | 6.51E-06 | 5.69E-03 | Downregulated |
| 0 | *Atp6v0b* | -0.5486 | 8.07E-05 | 5.64E-02 | Downregulated |
| 0 | *Magi3* | -0.8742 | 9.90E-05 | 5.77E-02 | Downregulated |
| 0 | *Prpf4b* | -0.474 | 1.57E-04 | 7.44E-02 | Downregulated |
| 0 | *Rbbp7* | -1.3129 | 1.91E-04 | 7.44E-02 | Downregulated |
| 0 | *Rims1* | -0.507 | 1.91E-04 | 7.44E-02 | Downregulated |
| 0 | *Dusp11* | -0.8453 | 2.35E-04 | 8.22E-02 | Downregulated |
| 1 | *Zfyve28* | 0.8092 | 1.12E-04 | 1.31E-01 | Upregulated |
| 1 | *Ptgds* | 1.0577 | 2.67E-04 | 1.87E-01 | Upregulated |
| 1 | *Rbbp7* | 1.4183 | 5.90E-04 | 2.06E-01 | Upregulated |
| 1 | *CT030170.6* | -0.9925 | 6.39E-06 | 2.24E-02 | Downregulated |
| 1 | *Ctnna2* | -0.596 | 5.62E-05 | 9.83E-02 | Downregulated |
| 1 | *Brsk1* | -1.1072 | 1.67E-04 | 1.46E-01 | Downregulated |
| 1 | *Ccl25* | -0.8767 | 3.63E-04 | 1.97E-01 | Downregulated |
| 1 | *Sppl3* | -0.974 | 4.13E-04 | 1.97E-01 | Downregulated |
| 1 | *Gabra2* | -1.0983 | 4.64E-04 | 1.97E-01 | Downregulated |
| 1 | *Stim1* | -0.7617 | 5.06E-04 | 1.97E-01 | Downregulated |
| 2 | *Cwc27* | 1.4362 | 1.25E-04 | 1.04E-01 | Upregulated |
| 2 | *Rcor1* | 1.511 | 7.22E-04 | 2.64E-01 | Upregulated |
| 2 | *Reep2* | -2.5474 | 2.11E-06 | 4.09E-03 | Downregulated |
| 2 | *Rbm28* | -1.7384 | 2.34E-06 | 4.09E-03 | Downregulated |
| 2 | *Ptprd* | -1.1693 | 7.23E-05 | 8.44E-02 | Downregulated |
| 2 | *Gpatch8* | -0.8352 | 1.49E-04 | 1.04E-01 | Downregulated |
| 2 | *Zc3h7b* | -1.9055 | 2.76E-04 | 1.61E-01 | Downregulated |
| 2 | *Ncam2* | -0.9158 | 4.85E-04 | 2.43E-01 | Downregulated |
| 2 | *Agrp* | -1.6572 | 7.45E-04 | 2.64E-01 | Downregulated |
| 2 | *Heatr3* | -1.7079 | 7.54E-04 | 2.64E-01 | Downregulated |
| 3 | *Adipor2* | 1.4047 | 7.44E-06 | 8.67E-03 | Upregulated |
| 3 | *Garnl3* | 2.3826 | 4.63E-05 | 2.70E-02 | Upregulated |
| 3 | *Fam171a1* | 2.5805 | 2.09E-04 | 9.12E-02 | Upregulated |
| 3 | *Sgk1* | 2.4287 | 3.91E-04 | 1.30E-01 | Upregulated |
| 3 | *Ppp1r13b* | 2.0932 | 3.94E-04 | 1.30E-01 | Upregulated |
| 3 | *Frmd4a* | -3.1281 | 4.96E-14 | 1.73E-10 | Downregulated |
| 3 | *9530059O14Rik* | -2.5915 | 2.02E-06 | 3.53E-03 | Downregulated |
| 3 | *Tnr* | -2.892 | 1.71E-05 | 1.49E-02 | Downregulated |
| 3 | *Abr* | -2.9181 | 4.44E-05 | 2.70E-02 | Downregulated |
| 3 | *Nfasc* | -1.1791 | 1.19E-04 | 5.94E-02 | Downregulated |

### Supplementary Table 10

| Cluster | Pathway name | Overlap | P value | Z-score |
|---|---|---|---|---|
| 0 | Synaptogenesis Signaling Pathway | 20/312 | 3.38E-09 | -2.828 |
| 0 | Opioid Signaling Pathway | 14/247 | 3.41E-06 | -0.535 |
| 0 | Synaptic Long Term Potentiation | 9/129 | 3.92E-05 | -1.414 |
| 0 | Phagosome Maturation | 9/151 | 1.33E-04 | N/A |
| 0 | nNOS Signaling in Skeletal Muscle Cells | 5/41 | 1.60E-04 | N/A |
| 1 | Huntington's Disease Signaling | 7/239 | 5.26E-03 | N/A |
| 1 | Dolichol and Dolichyl Phosphate Biosynthesis | 1/2 | 1.75E-02 | N/A |
| 1 | **β**-alanine Degradation I | 1/2 | 1.75E-02 | N/A |
| 1 | Glutamine Degradation I | 1/2 | 1.75E-02 | N/A |
| 1 | Pyridoxal 5'-phosphate Salvage Pathway | 3/66 | 2.03E-02 | N/A |
| 2 | Synaptogenesis Signaling Pathway | 11/312 | 8.74E-04 | -2.530 |
| 2 | G Beta Gamma Signaling | 6/122 | 2.62E-03 | -1.633 |
| 2 | Ephrin Receptor Signaling | 7/189 | 5.73E-03 | -1.342 |
| 2 | Netrin Signaling | 4/65 | 6.22E-03 | -2.000 |
| 2 | Neuregulin Signaling | 5/105 | 6.76E-03 | -0.447 |
| 3 | Fc**γ** Receptor-mediated Phagocytosis in Macrophages and Monocytes | 5/94 | 3.59E-03 | -0.447 |
| 3 | RhoGDI Signaling | 7/189 | 4.63E-03 | 0.816 |
| 3 | Huntington's Disease Signaling | 8/239 | 4.66E-03 | 2.000 |
| 3 | Remodeling of Epithelial Adherens Junctions | 4/68 | 6.35E-03 | N/A |
| 3 | ERK5 Signaling | 4/72 | 7.76E-03 | 0.000 |

N/A = Not available
