## Supplemental Figures for "A survey of the mouse hindbrain in the fed and fasted state using single-nucleus RNA sequencing"

### Supplementary Figure 1

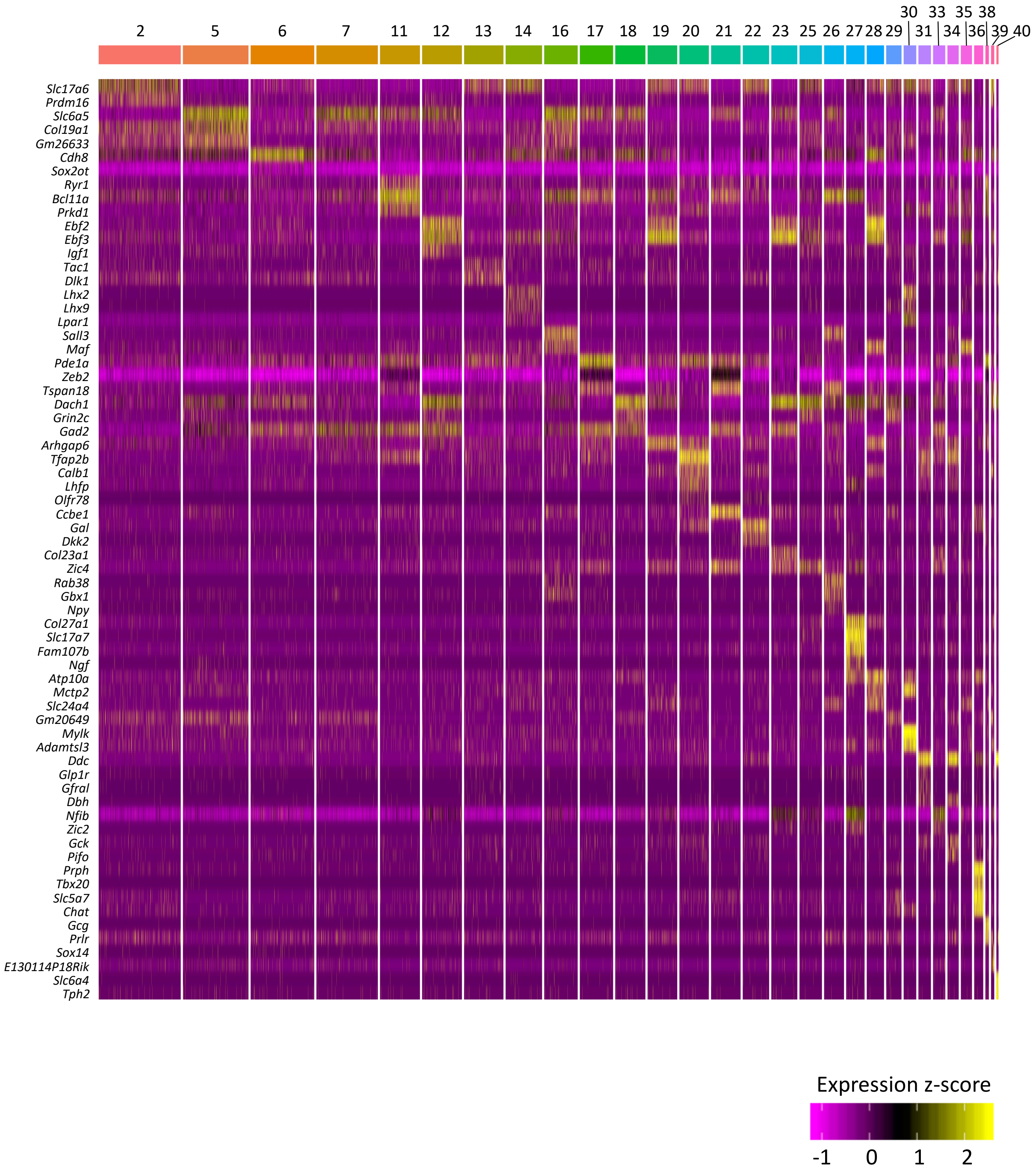

### Supplementary figure 2

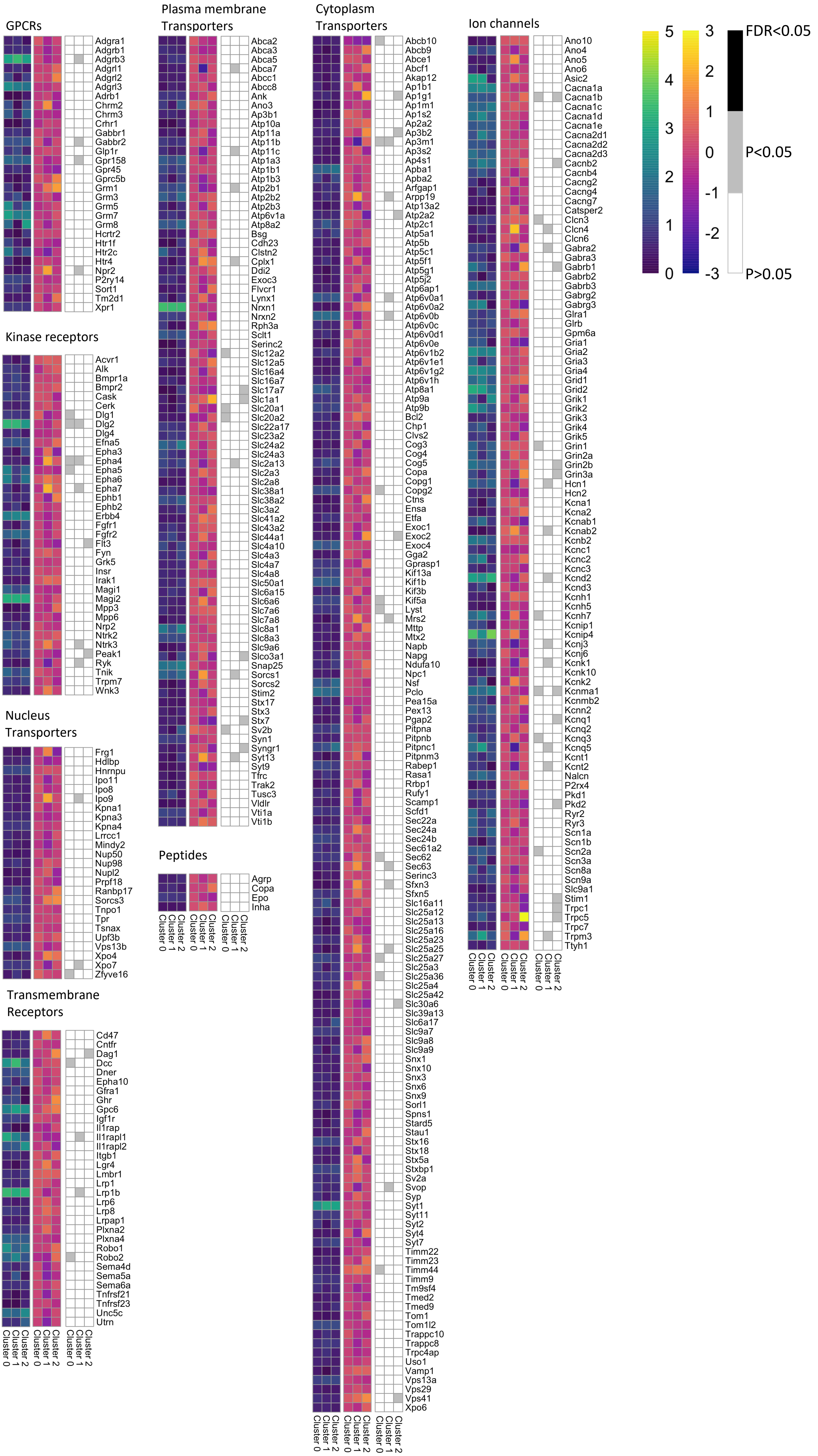

### Supplementary figure 3

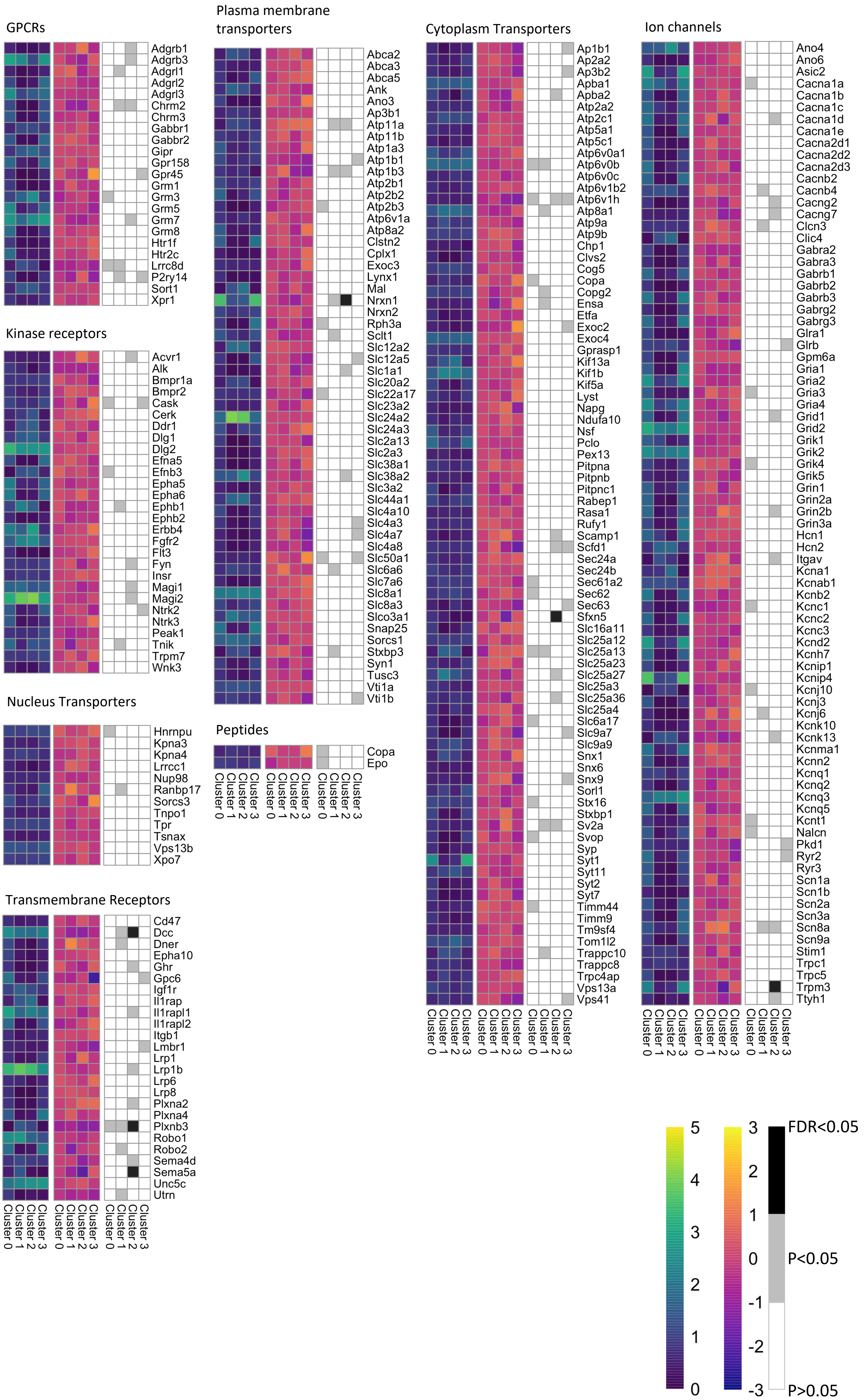

### Supplementary figure 4

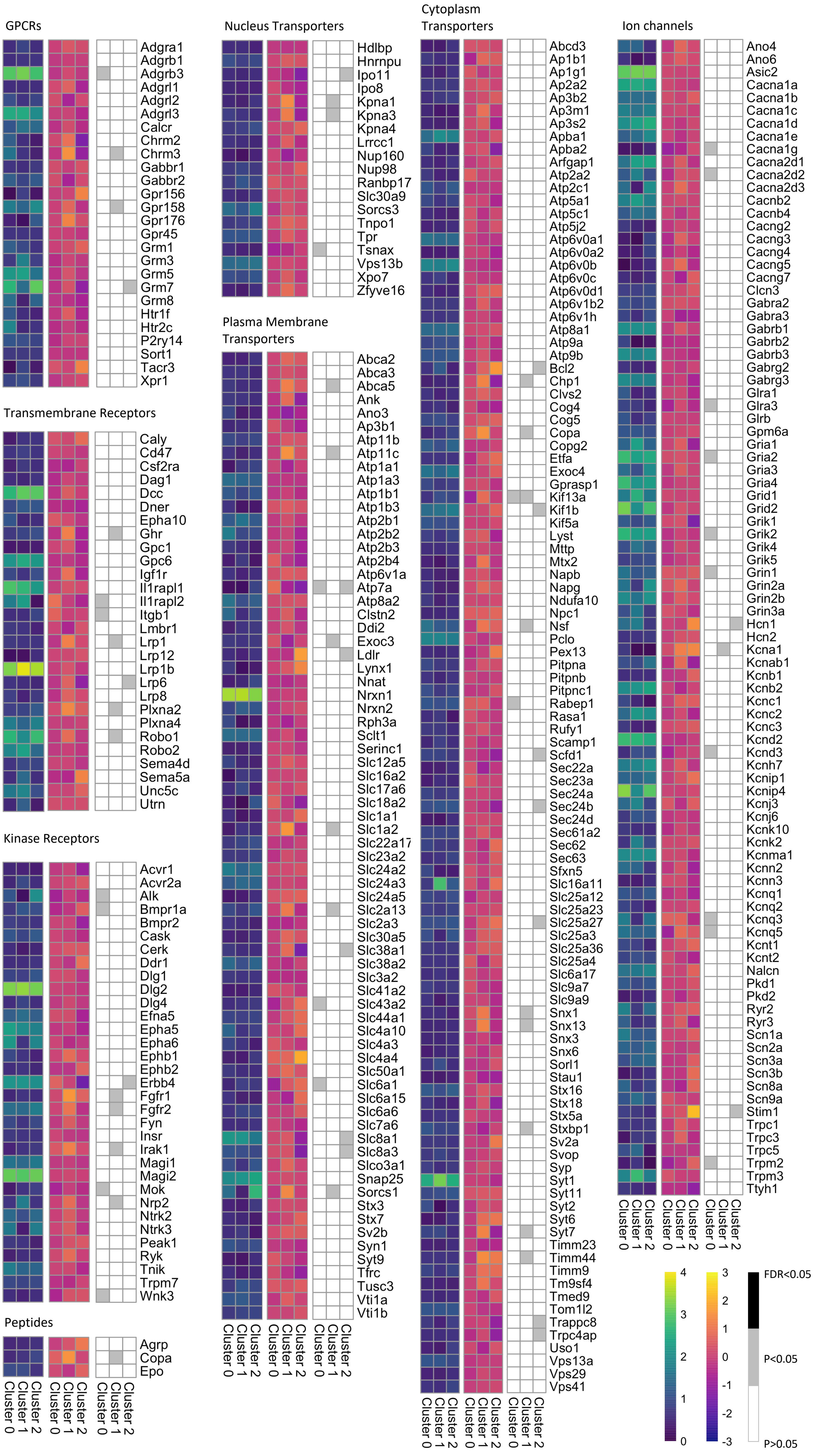

### Supplementary figure 5

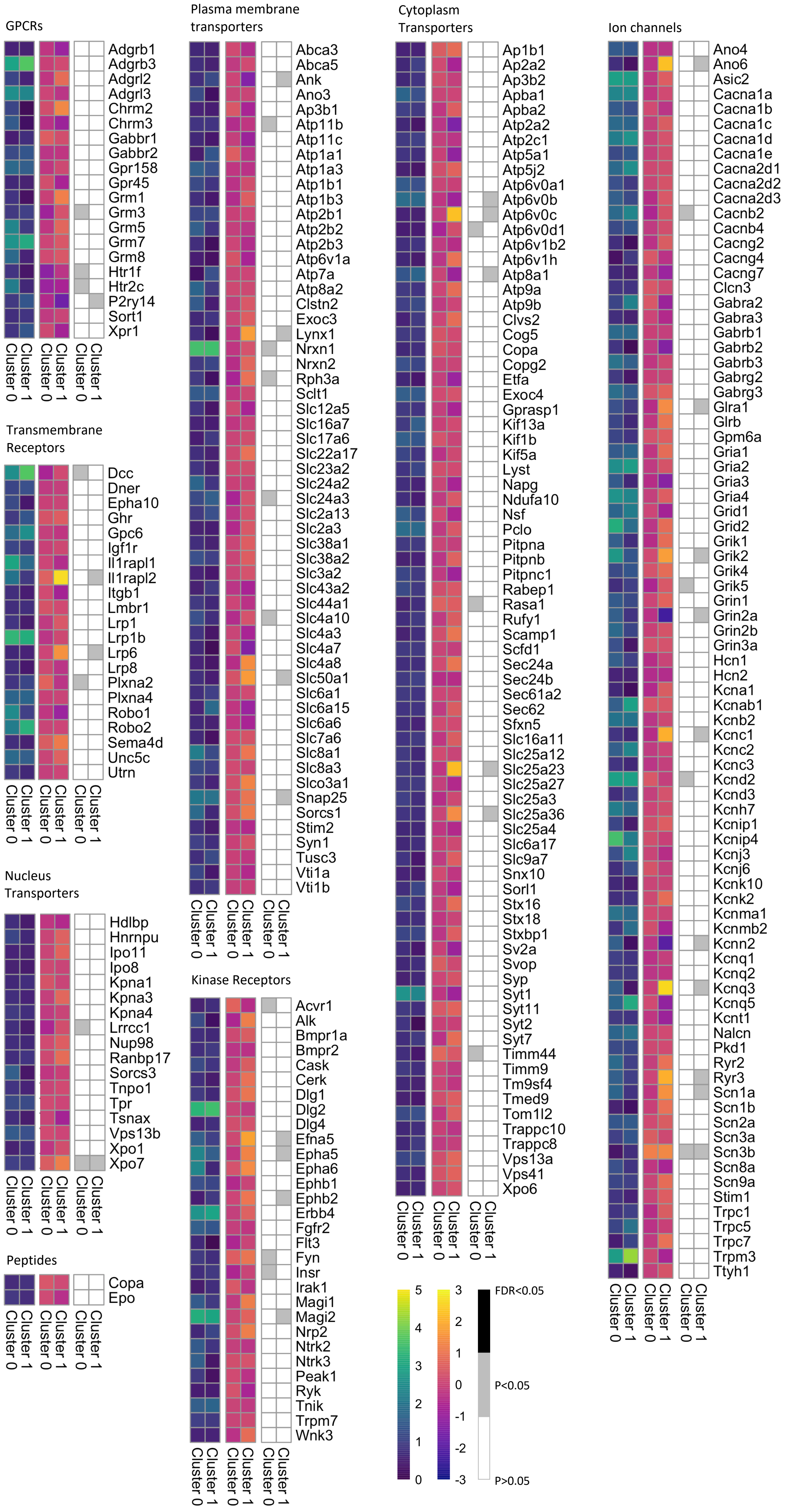

### Supplementary figure 6

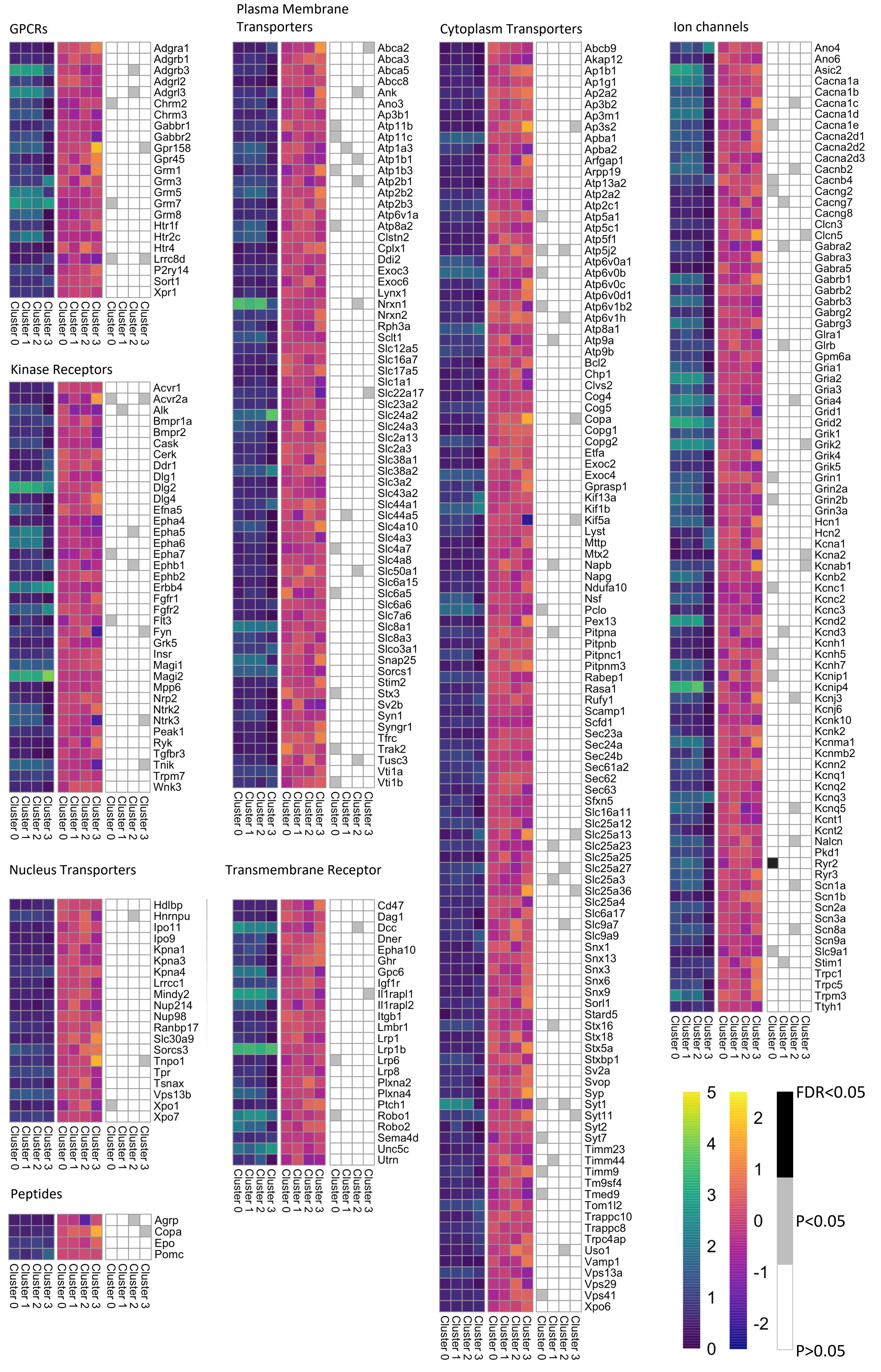
